# *PSEUDOMONAS AERUGINOSA* VOLATILOME CHARACTERISTICS AND ADAPTATIONS IN CHRONIC CYSTIC FIBROSIS LUNG INFECTIONS

**DOI:** 10.1101/2020.06.13.126698

**Authors:** Trenton J. Davis, Ava V. Karanjia, Charity N. Bhebhe, Sarah B. West, Matthew Richardson, Heather D. Bean

## Abstract

*Pseudomonas aeruginosa* chronic lung infections in individuals with cystic fibrosis (CF) significantly reduce quality of life and increase morbidity and mortality. Tracking these infections is critical for monitoring patient health and informing treatments. We are working toward the development of novel breath-based biomarkers to track chronic *P. aeruginosa* lung infections *in situ*. Using comprehensive two-dimensional gas chromatography coupled to time-of-flight mass spectrometry (GC×GC-TOFMS), we characterized the *in vitro* volatile metabolomes (or volatilomes) of 81 *P. aeruginosa* isolates collected from 17 CF patients over at least a five-year period of their chronic lung infections. We detected 539 volatiles produced by the *P. aeruginosa* isolates, 69 of which were core volatiles that were highly conserved. We found that each early infection isolate has a unique volatilome, and as infection progresses, the volatilomes of isolates from the same patient become increasingly dissimilar, to the point that these intra-patient isolates are no more similar to one another than to isolates from other patients. We observed that the size and chemical diversity of *P. aeruginosa* volatilomes do not change over the course of chronic infections; however, the relative abundances of core hydrocarbons, alcohols, and aldehydes do change, and are correlated to changes in phenotypes associated with chronic infections. This study indicates that it may be feasible to track *P. aeruginosa* chronic lung infections by measuring changes to the infection volatilome, and lays the groundwork for exploring the translatability of this approach to direct measurement using patient breath.

**IMPORTANCE:** *Pseudomonas aeruginosa* is a leading cause of chronic lung infections in cystic fibrosis (CF), and are correlated with lung function declines. Significant clinical efforts are, therefore, aimed at detecting infections and tracking them for phenotypic changes, such as mucoidy and antibiotic resistance. Both the detection and tracking of lung infections relies on sputum cultures, but due to improvements in CF therapies, sputum production is declining though risks for lung infections persist. Therefore, we are working toward the development of breath-based diagnostics for CF lung infections. In this study we characterized of the volatile metabolomes of 81 *P. aeruginosa* clinical isolates collected from 17 CF patients over a duration of at least five years of a chronic lung infection. We found that the volatilome of *P. aeruginosa* adapts over time, and correlates to infection phenotype changes, suggesting it may be possible to track chronic CF lung infections with a breath test.

## INTRODUCTION

Cystic fibrosis (CF) is an autosomal recessive disease caused by a mutation in the CFTR protein that regulates ion transport across epithelia. In the lungs, reduced or lost CFTR function leads to defective mucociliary transport, facilitating infection and colonization by a plethora of microorganisms (1). *Pseudomonas aeruginosa* is one of the most prevalent lung pathogens in CF – especially after adolescence (2) – and is able to establish chronic infections that can last for years to decades (3, 4). *P. aeruginosa* lung infection is associated with more rapid lung function decline, increased risk of hospitalization, and increased risk of death (5, 6). Significant clinical efforts are, therefore, aimed at diagnosing and treating new infections to delay the onset of chronic infection (7).

As *P. aeruginosa* transitions from an acute infection to chronicity, it undergoes a variety of genotypic, phenotypic, and metabolic changes (8, 9). It has been well-established that several phenotypes are correlated with chronic infection, including reduced motility and increased mucoidy, antibiotic resistance, and biofilm formation (9-11), and some of these phenotypes are also correlated with poorer patient outcomes (6, 12, 13). In light of this, *P. aeruginosa* phenotypes could be used to evaluate infection stage, CF disease progression, and patient morbidity risk (10, 12, 14, 15). Poor access to infection sites in the lower airways, however, delays the detection of new *P. aeruginosa* infections and reduces the feasibility and accuracy of disease state tracking via bacterial phenotypes, whether by culture-dependent or culture-independent methods (16).

We are developing breath-based diagnostics to detect new *P. aeruginosa* infections and track chronic infection phenotypes in the CF lung. Breath consists of thousands of volatile organic compounds containing a wealth of information about human health and disease (17, 18), and is being leveraged for the development of diagnostics for wide-ranging conditions (19, 20). In the context of chronic lung infections, breath-based diagnostics possess significant advantages over other diagnostic modalities. First, breath provides a non-invasive way of sampling the entire ventilated lung, a limitation of both sputum and bronchoalveolar lavage fluid that leads to delayed diagnosis of new infections, or the detection of infection phenotype changes (16, 21-24). Second, breath sampling captures metabolic information about the pathogens *in situ*, eliminating the need for microbial enrichment steps and speeding the time-to-diagnosis by days to weeks (25, 26). Additionally, *in situ* metabolic measurements provide information on the physiology of the pathogens in the context of disease versus the context of the lab. We propose that a breath-based diagnostic can be created for detecting and tracking CF *P. aeruginosa* lung infections by correlating the microbial genotypic and phenotypic changes that occur during chronic infections to changes in the infection volatile metabolome, or “volatilome”.

In order to develop a set of biomarkers for diagnosing and tracking *P. aeruginosa* lung infections in CF, we must consider the genomic, phenomic, and volatilomic diversity of the species, and how that diversity may be altered via adaptation to the CF lung environment. A recent effort to sequence more than 1300 *P. aeruginosa* isolates demonstrates that the genomes are highly flexible, with only 1% of the pan-genome conserved across all isolates (27). An untargeted analysis of the volatile metabolites produced *in vitro* by 24 *P. aeruginosa* clinical isolates shows that the volatilomes are also diverse, with 18% of the pan-volatilome conserved in this comparatively small study (28). In the context of CF, most early *P. aeruginosa* infections in young patients are caused by unique strains that come from the patients’ environments (29-32). We therefore expect the volatilomes early in infection to have high inter-patient dissimilarity, reflecting the genomic diversity of *P. aeruginosa* CF isolates (3, 33-35), and to be chemically diverse (*i*.*e*., volatiles from a wide range of chemical classes). Once chronic infections are established, however, the “founder” *P. aeruginosa* strain evolves into a population of clonal sub-strains (3, 8, 36-40), many of which harbor CF-typical loss-of-function mutations in regulatory genes and sigma factors (*e*.*g*., *lasR* and *mucA*) (3, 41, 42). From this, we posit that the size of the *P. aeruginosa* volatilome will shrink during chronic infection, with a concomitant decrease in chemical diversity and a reduction in inter-patient dissimilarity.

The primary goal of this study is to build a foundation for using volatilomes to diagnose and track *P. aeruginosa* CF lung infections by exploring how the *in vitro* volatilome of *P. aeruginosa* CF isolates change over the course of chronic infection. We used comprehensive two-dimensional gas chromatography coupled with time-of-flight mass spectrometry (GC×GC-TOFMS) to characterize the volatilomes of 81 *P. aeruginosa* isolates from early and late chronic lung infections from 17 persons with CF. Using an untargeted metabolomics approach, we characterized the size and chemical composition of the CF *P. aeruginosa* volatilome, and identified core volatiles that would be a primary source of breath biomarkers for diagnosing infections. We also investigated how the *P. aeruginosa* volatilome is shaped by the CF lung environment, characterizing changes in volatilome sizes and compositions over time, and how these changes relate to intra-patient and inter-patient volatilome dissimilarities. A secondary goal of this study is to provide further data on the *P. aeruginosa* volatilome via the largest single analysis of *P. aeruginosa* headspace volatiles, to-date.

## METHODS

### Bacterial isolates

Eighty one *P. aeruginosa* isolates from 17 individuals with CF were acquired from the Cystic Fibrosis Isolate Core at Seattle Children’s Center for Global Infectious Disease Research. From the majority of patients we obtained three isolates: one early isolate, defined as the first cultured *P. aeruginosa* isolate, and two late isolates collected a minimum of five years after the first isolate. For one patient (Patient 23), 32 additional isolates were collected at intervals over 7.5 years between the collections of the early and two late isolates. For one patient, only one isolate was included in this study. The full strain names of the isolates are provided in Table S1, and all isolates are available upon request to the CF Isolate Core (https://www.seattlechildrens.org/research/resources/cystic-fibrosis-isolate/).

We quantified five phenotypes that are correlated with chronic infections: proteases, pyocyanin, rhamnolipids, twitching motility, and mucoidy. *P. aeruginosa* was cultured from glycerol stocks on LB agar for 24 h, then a single colony was cultured aerobically to stationary phase in LB Lennox broth (10 g tryptone, 5 g yeast extract, 5 g NaCl per liter) at 37 °C with shaking at 200 RPM, unless otherwise noted. Results of phenotype assays are reported in Table S1 as means of biological triplicates, except for mucoidy for which five replicates were measured. Pyocyanin production was evaluated by methods adapted from (43). Cell-free *P. aeruginosa* culture supernatant (7.5 mL) was extracted with 4.5 mL of chloroform, inverted for 2 min and centrifuged at 4,122 × *g* for 15 min. Three milliliters of the organic phase was removed and extracted with 750 µL of 0.2 *N* hydrochloric acid and vortexed for 2 min. The aqueous phase was aliquoted into a 96-well plate and absorbance was measured at 520 nm. Protease production was evaluated by methods adapted from (15). A sterile wooden inoculation stick was dipped into a culture of *P. aeruginosa* and then gently touched to the surface of a brain-heart infusion-skim milk agar (1.5%) plate. Plates were incubated upright at 37 °C and zones of clearance were measured at 48 h. Rhamnolipids production was evaluated by methods adapted from (44). Proteose peptone-glucose-ammonium salts (PPGAS) medium was inoculated from an LB pre-culture of *P. aeruginosa* and cultured at 37 °C to stationary phase with shaking at 200 RPM. Cultures were centrifuged at 21,694 × *g* for 1 min to pellet cells. The supernatant was serially diluted two-fold in PPGAS and 20 µL of each dilution was spotted onto a microtiter plate lid. Spots were classified as drops or collapsed drops and assigned numeric scores corresponding to the number of dilutions needed to obtain a drop. Twitching motility was evaluated by methods adapted from (45, 46). The pointed end of a sterile toothpick was touched to the edge of a single *P. aeruginosa* colony and then stabbed to the bottom of a twitching motility agar plate (per 1 L: 10 g tryptone, 5 g NaCl, 5 g yeast extract, 10 g agar). Plates were incubated at 37 °C for 24 h and the radius of the interstitial biofilm was measured. For mucoidy, frozen glycerol stocks of *P. aeruginosa* were streaked onto LB Lennox agar (1.5%) plates and grown for 48 hrs. Colonies were visually inspected and the degree of mucoid morphology was scored from 0 to 2, in 0.5 increments (0 = highly mucoid, 2 = non-mucoid). Twenty pairs of isolates (one Early and one Late) from ten patients were selected for additional statistical analyses, described below, based on changes to their phenotypes during chronic infections.

### Sample preparation

Isolates were cultured as previously described (47). Briefly, isolates were cultured aerobically for 16 h at 37 °C in 5 mL of lysogeny broth-Lennox (LB), and then diluted 1000-fold into 25 mL of fresh LB and grown for 24 h under the same conditions. For metabolomics analyses, cells were removed via centrifugation through a 0.2 *µ*m filter and 2 mL of each filtrate was transferred to a 10 mL GC headspace vial with screw caps. Samples were prepared in biological triplicate, with LB media controls prepared in parallel, and stored at –20 °C prior to analysis.

### GC×GC analysis and data processing

Culture filtrates were thawed and maintained at 4 °C until analyzed, as previously described (47). Headspace volatiles were characterized using a Pegasus® 4D GC×GC-TOFMS (LECO® Corporation, St. Joseph, MI) equipped with a MPS Pro® rail autosampler (Gerstel® Inc, Linthicum Heights, MD). Column set configuration, and GC×GC, MS, and data processing method parameters are previously reported (47), and summarized in Miscellaneous Table 1 (https://doi.org/10.1101/2020.06.13.126698).

**Table 1.** Putatively identified volatile compounds metabolized by *P. aeruginosa* CF isolates.

| Compound Name<sup>a</sup> | Formula | Chemical Class | Samples ($n = 81$) | Refs.<sup>b</sup> |
| --- | --- | --- | --- | --- |
| **Core volatile compounds** |  |  |  |  |
| **3-methylbutanal** | C<sub>5</sub>H<sub>10</sub>O | Aldehyde | 79 |  |
| thiocyanic acid, methyl ester | C<sub>2</sub>H<sub>3</sub>NS | Ester | 77 | (53) |
| 3-methyl-1-butanol | C<sub>5</sub>H<sub>12</sub>O | Alcohol | 77 | (58, 60, 61) |
| **pyridine** | C<sub>5</sub>H<sub>5</sub>N | Aromatic | 77 |  |
| **3-penten-2-one** | C<sub>5</sub>H<sub>8</sub>O | Ketone | 77 |  |
| **3-methyl-1-buten-1-ol** | C<sub>5</sub>H<sub>10</sub>O | Alcohol | 78 |  |
| 4-methyl-3-penten-2-one | C<sub>6</sub>H<sub>10</sub>O | Ketone | 81 | (28, 62) |
| hexanal | C<sub>6</sub>H<sub>12</sub>O | Aldehyde | 81 | (58, 60, 62) |
| **heptanal** | C<sub>7</sub>H<sub>14</sub>O | Aldehyde | 80 |  |
| 2,5-dimethylpyrazine | C<sub>6</sub>H<sub>8</sub>N<sub>2</sub> | Aromatic | 81 | (53) |
| 2-nonanone | C<sub>9</sub>H<sub>18</sub>O | Ketone | 81 | (28, 53, 58, 59, 61-67) |
| **4-ethyl-1,2-dimethylbenzene** | C<sub>10</sub>H<sub>14</sub> | Aromatic | 77 |  |
| **1,3,3-trimethyl-(bicyclo)-heptan-2-ol** | C<sub>10</sub>H<sub>18</sub>O | Alcohol | 78 |  |
| **1-(4-ethylphenyl)-ethanone** | C<sub>10</sub>H<sub>12</sub>O | Ketone | 79 |  |
| **Non-core volatile compounds** |  |  |  |  |
| dimethyl sulfide | C<sub>2</sub>H<sub>6</sub>S | Other | 29 | (28, 53, 58, 60, 61, 64, 68, 69) |
| **2-methyl-3-buten-2-ol** | C<sub>5</sub>H<sub>10</sub>O | Alcohol | 35 |  |
| 2-butanone | C<sub>4</sub>H<sub>8</sub>O | Ketone | 1 | (28, 53, 58, 59, 61, 63, 66, 70) |
| **tetrahydrofuran** | C<sub>4</sub>H<sub>8</sub>O | Aromatic | 1 |  |
| **acetic acid** | C<sub>2</sub>H<sub>4</sub>O<sub>2</sub> | Carboxylic acid | 55 |  |
| methyl thioacetate | C<sub>3</sub>H<sub>6</sub>OS | Thiol | 3 | (58, 60, 62) |
| 2,4-dimethylfuran | C<sub>6</sub>H<sub>8</sub>O | Aromatic | 66 | (28) |
| **butanoic acid, methyl ester** | C<sub>5</sub>H<sub>10</sub>O<sub>2</sub> | Ester | 3 |  |
| **3-hydroxybutan-2-one** | C<sub>4</sub>H<sub>8</sub>O<sub>2</sub> | Ketone | 18 |  |
| dimethyl disulfide | C<sub>2</sub>H<sub>2</sub>S<sub>2</sub> | Other | 1 | (28, 58-62, 66, 68-73) |
| **2-methyl-butanoic acid, methyl ester** | C<sub>6</sub>H<sub>12</sub>O<sub>2</sub> | Ester | 3 |  |
| **3-methyl-butanoic acid, methyl ester** | C<sub>6</sub>H<sub>12</sub>O<sub>2</sub> | Ester | 3 |  |
| furfural | C<sub>5</sub>H<sub>4</sub>O<sub>2</sub> | Aromatic | 67 | (60) |
| **2-butylfuran** | C<sub>8</sub>H<sub>12</sub>O | Aromatic | 6 |  |
| 2-hexen-1-ol | C<sub>6</sub>H<sub>12</sub>O | Alcohol | 51 | (28) |
| **1-butoxy-2-propanol** | C<sub>7</sub>H<sub>16</sub>O<sub>2</sub> | Alcohol | 61 |  |
| **6-methyl-2-heptanone** | C<sub>8</sub>H<sub>16</sub>O | Ketone | 33 |  |
| **1-methylethylbenzene** | C<sub>9</sub>H<sub>12</sub> | Aromatic | 57 |  |
| 3-octanone | C<sub>8</sub>H<sub>16</sub>O | Ketone | 56 | (28, 61) |
| **4,6-dimethyl-2-heptanone** | C<sub>9</sub>H<sub>18</sub>O | Ketone | 47 |  |
| **2-ethyl-5-methylpyrazine** | C<sub>7</sub>H<sub>10</sub>N<sub>2</sub> | Aromatic | 58 |  |
| 4-nonanone | C<sub>9</sub>H<sub>18</sub>O | Ketone | 49 | (28) |
| **tetramethylpyrazine** | C<sub>8</sub>H<sub>12</sub>N<sub>2</sub> | Aromatic | 5 |  |
| 2-octenal | C<sub>8</sub>H<sub>14</sub>O | Aldehyde | 48 | (53) |
| octanenitrile | C<sub>8</sub>H<sub>15</sub>N | Other | 73 | (28) |
| 2,4-octadienal | C<sub>8</sub>H<sub>12</sub>O | Aldehyde | 9 | (28) |
| 3-decanone | C<sub>10</sub>H<sub>20</sub>O | Ketone | 41 | (53) |
| **decanal** | C<sub>10</sub>H<sub>20</sub>O | Aldehyde | 65 |  |
| 2-decanone | C<sub>10</sub>H<sub>20</sub>O | Ketone | 13 | (28) |
| **4-undecanone** | C<sub>11</sub>H<sub>22</sub>O | Ketone | 30 |  |
| 2-dodecanone | C<sub>12</sub>H<sub>24</sub>O | Ketone | 18 | (28, 53) |
| **2-butyl-1-octanol** | C<sub>12</sub>H<sub>26</sub>O | Alcohol | 48 |  |
| **2,4-bis(1,1-dimethyl)-phenol** | C<sub>14</sub>H<sub>22</sub>O | Aromatic | 26 |  |

<sup>a</sup>Bolded compounds have not been previously reported as *P. aeruginosa* volatile compounds.
<sup>b</sup>Previous references for these volatiles in *P. aeruginosa* cultures are provided.

Data processing steps are outlined in Miscellaneous Figure 1. Chromatographic artifacts and suspected contaminants based on peak names and comparisons to blanks were removed (Miscellaneous Table 2), as well as poorly modulated peaks eluting prior to 358 s (acetone retention time) in the first dimension. Missing values were handled as follows: if detected in one of three replicates, the measured value was permuted to 0; if detected in two of three replicates, the missing value was imputed as half of the minimum detected value for that compound across all samples.

**FIG 1.**
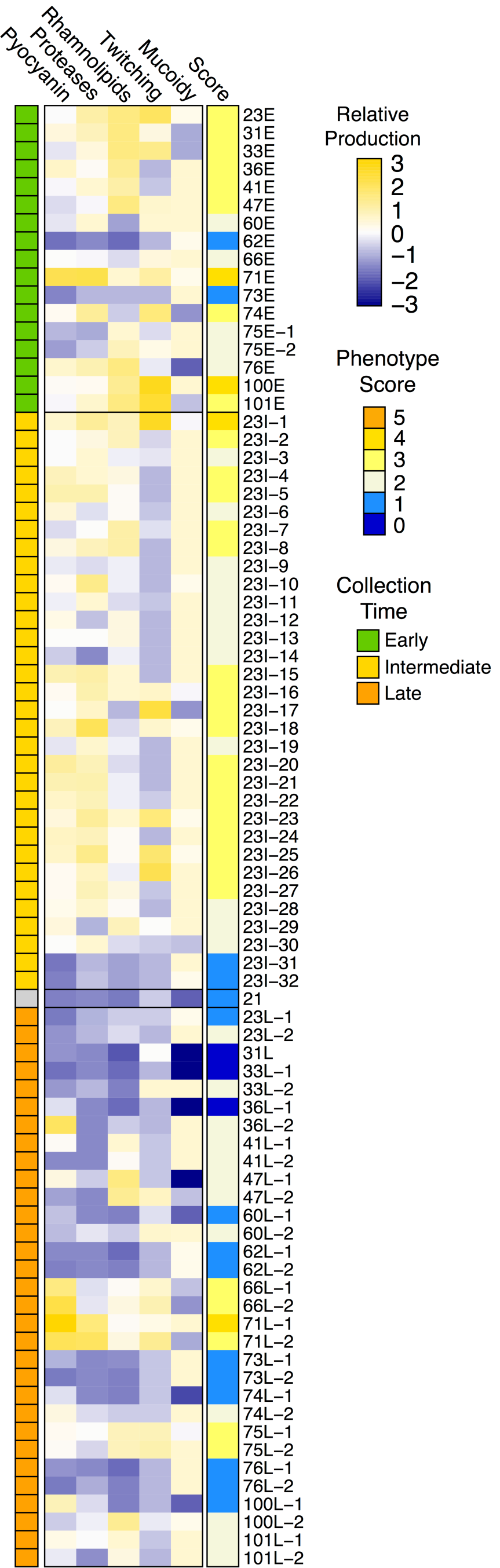
Relative phenotype production of 81 *P. aeruginosa* CF chronic lung infection isolates. Each phenotype was independently scaled to a range of 0 to 1, with 0 representing the minimum data point and 1 representing the maximum data point (excluding outliers), and the relative productions are depicted in the heat map as standardized values (mean-centered and scaled to unit variance) across isolates. Phenotype Score (“Score”) is the sum of the scaled, non-standardized phenotype values, rounded to the nearest integer.

Probabilistic quotient normalization (PQN) was applied to account for differences in peak abundance due to variations in culture cell density, followed by a log_10_ transformation (48). Peaks were further filtered out based on the following criteria: (1) within-triplicate intraclass correlation coefficient (ICC) < 0.75 (definition: absolute agreement; model: two-way mixed effects; type: mean of *k* measurements) (49); and (2) only detected in sterile media, or not significantly greater in abundance compared to sterile media using the Wilcoxon rank-sum test with Benjamini-Hochberg adjustment (significance threshold of .05) (50).

For statistical analyses beyond the reporting of detected peaks, peaks that showed significant correlations to run order were identified using Kendall’s tau with Benjamini-Hochberg adjustment (−0.6 ≤ τ ≥ 0.6, significance threshold of .05) and removed. Principal component analysis (PCA) revealed an apparent batch effect that was attributed to, at least in part, a non-biological phenomenon and was described previously (47). This batch effect was corrected using an empirical Bayes approach (51). The geometric means of triplicates of the batch-corrected data were used for further statistical analyses. For analysis of the ten paired early and late chronic infection isolates, batch-corrected data was not used as all of these isolates were contained within the same batch.

Peaks were putatively identified using published reporting standards (52). Level 2 identifications were determined based on the following criteria: (1) ≥ 80% mass spectral forward match using the NIST 2011 library; (2a) first dimension retention times that possess a strong linear fit with carbon number in homologous series (*r*^2^ ≥ 0.995, for the identification of select 2-, 3-, and 4-ketones), or (2b) experimentally determined linear retention indices (LRI) consistent with published LRIs, as determined by the following acceptance criteria based on the differences between experimental and published non-polar and polar column RIs for Grob’s test

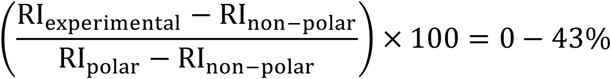

Level 3 identifications were based on at least an 80% mass spectral forward match score. Chemical classifications (alcohols, aldehydes, aromatics, carboxylic acids, esters, hydrocarbons, ketones, thiols, or other) were assigned based primarily on their mass spectral identity, and secondarily by their chromatographic characteristics. Using polar second dimension columns, compounds of differing functional classes elute at easily discernible second-dimension retention times in a stratified manner, typically (from smallest ^2^*t*_R_ to largest): hydrocarbons < ketones < aromatics (53-57). In this study, compounds identified as hydrocarbon, ketones, and aromatics eluted at approximately 0.71 s ± 0.05 s, 0.88 s ± 0.22 s, and 1.12 s ± 0.26 s, respectively. Level 4 identifications are those that meet none of the above-mentioned criteria.

### Statistical analyses

A Phenotype Score was determined, for each replicate, by scaling the data for each phenotype to a range of 0 to 1, where 1 is the maximum value and 0 is the minimum value (excluding outliers). Scaled phenotype data were then summed to yield the Score, and averaged across replicates. Patient age was not included in calculation of Scores. Two-tailed Wilcoxon signed-rank tests (with continuity correction, where appropriate; significant threshold of .05) were used to test for significant differences between Late and Early isolate Scores.

Volatile compounds that were detected in at least 95% of all samples were classified as “core” volatiles. Significant differences between inter-isolate phenotypes were tested using the Wilcoxon signed-rank test (significance threshold of .05). Significant differences between intra- and inter-patient volatilomes were tested using one-way analysis of variance (ANOVA) (significance threshold of .05), and Tukey’s honestly significant differences (HSD) multiple comparison procedure (significance threshold of .05). The relatedness of isolates based on their volatilomes were assessed using agglomerative hierarchical clustering analysis (HCA), non-metric multidimensional scaling (NMDS), and permutational multivariate analysis of variances (PERMANOVA) on the Euclidean distances between isolates. Linear regression and Pearson’s correlation of Euclidean distances of Patient 23 isolates were used to assess intra-patient isolate dissimilarity over time. All statistical analyses were performed using R version 3.3.2 (The R Foundation for Statistical Computing) with the following packages (version): *ICC* (2.3.0), *pairwiseAdonis* (0.0.1), *pheatmap* (1.0.8), *stats* (3.3.2), and *sva* (3.22.0).

## Data availability

Metabolomic data (chemical feature peak areas and retention time information) included in this study are available at the NIH Common Fund’s National Metabolomics Data Repository (NMDR) website, the Metabolomics Workbench, at www.metabolomicsworkbench.org, where it has been assigned Project ID PR000970, Study ID ST001414 (DOI: http://dx.doi.org/10.21228/M89Q4F). Additional Miscellaneous Information mentioned can be found alongside Supplementary Information at bioRxiv (DOI: https://doi.org/10.1101/2020.06.13.126698)

## RESULTS

### Characteristics of *P. aeruginosa* CF isolates

We obtained from a biorepository 81 *P. aeruginosa* chronic infection isolates, which had been collected from 17 individuals with CF. From 14 patients, two or three *P. aeruginosa* isolates were obtained: one isolate that was the first cultured *P. aeruginosa*, and one or two isolates that were collected at least five years after the first. We refer to these isolates as Early and Late, respectively. For one patient (Patient 75), four isolates were collected: two Early isolates collected one month apart and two Late isolates, collected ten and 16 years after the first. For one patient (Patient 23), 32 additional isolates were collected over the course of a 7.5 y infection period, which we have termed Intermediate isolates. For one patient, only one isolate was collected. These isolates were genetically characterized in a study by Smith *et al*., who determined that the intra-patient replicate isolates are all clonally related, with the exception of the four isolates from patient 75, which are actually two clonally-related Early/Late pairs (isolates 75E-1 and 75L-2 are clonal, and isolates 75E-2 and 75L-1 are clonal) (3).

For all isolates, we characterized five clinically relevant phenotypes *in vitro*: mucoidy, pyocyanin production, rhamnolipids production, protease production, and twitching motility (Fig. 1; Table S1). These phenotypes are commonly altered during the course of chronic CF lung infections (15). There were wide ranges in the expression of these phenotypes across all isolates, as expected. Several isolate sets exhibited phenotypes consistent with age of collection in that Early isolates possessed higher degrees of motility, higher quantities of pyocyanin, rhamnolipids, and proteases (indicating intact quorum regulation), and lower mucoidy compared to their cognate Late isolates (*e*.*g*., Patients 23, 31, 33, 36, and 76). The associations between relative patient age at isolate collection and *P. aeruginosa* phenotypes were not perfect, however. For example, Early isolate 62E had no detectable pyocyanin, proteases, rhamnolipids, or twitching motility, and many of the Late isolates retained more early-like phenotypes (e.g., both Late isolates of 41, 66, 71, 75, and 101, and 60L-1, 74L-2, and 100L-2). Despite observable changes in isolate phenotype within patient, no phenotypes were significantly different when comparing Late isolate(s) to the Early isolate. Collectively, the 81 isolates were highly varied, and represented the array of phenotypes we expect to observe in the span of *P. aeruginosa* CF lung infections from initial infection to long-established chronic infections.

### The *in vitro* volatilome of *P. aeruginosa* CF isolates

Taking an untargeted metabolomics approach, we cultured the clinical *P. aeruginosa* isolates *in vitro* and characterized their volatilomes using GC×GC-TOFMS. Following extensive data processing, including removal of analytically biased chromatographic features, we conservatively attribute 539 non-redundant volatile compounds to the growth and metabolism of the *P. aeruginosa* isolates (Table S2). Among these, 69 compounds are shared by at least 95% (*n* ≥ 77) of the isolates, representing core volatiles. Using minimum metabolomics reporting standards, we assigned compound identification levels 1-4 (1 high, with 2 being the highest in this study) based on a combination of mass spectral and chromatographic characteristics. We were able to assign putative names to 14 core and 33 non-core compounds with an ID level 2 (Table 1), many of which have been previously reported as *P. aeruginosa* volatiles (Table 1). While we detected 2-aminoacetophenone (the volatile compound responsible for the grape-like odor of *P. aeruginosa*), it possessed an intraclass correlation slightly lower than the applied threshold of 0.75 and was ultimately filtered out of our peak list. We also identified several compounds of a variety of chemical classes that have not been previously reported for *P. aeruginosa*, including: alcohols (3-methyl-1-buten-1-ol, 1-butoxy-2-propanol, 2-butyl-1-octanol), esters (butanoic acids, methyl isovalerate), aromatics (tetrahydrofuran, tetramethylpyrazine, 2-butylfuran, 3-cyano-2,5-dimethylpyrazine, 2,4-(1,1-dimethyl)-phenol), unsaturated ketones (3-hydroxy-butan-2-one, 4-undecanone, 1-octen-3-one, 3-penten-2-one, 6-methyl-2-heptanone, 1- (4-ethylphenyl)-ethanone), and an aldehyde (decanal).

Chemical classifications were assigned to ID level 2 and 3 compounds (187 compounds) using mass spectral match and retention time characteristics (Fig. 2). Among Level 2 and 3 core volatiles, the majority of classified compounds were hydrocarbons (41%), followed by ketones (16%). Other oxidized compounds, including alcohols, aromatics, acids, and thiols, accounted for an additional 43%. The non-core volatiles had similar chemical compositions, which are consistent with previous studies of the *P. aeruginosa* volatilome (25, 28, 53, 58, 59). Of the entire volatilome, 65% of the volatiles (*n* = 352) possessed less than an 80% mass spectral match to the 2011 NIST MS library, and as such were classified as unknowns.

**FIG 2.**
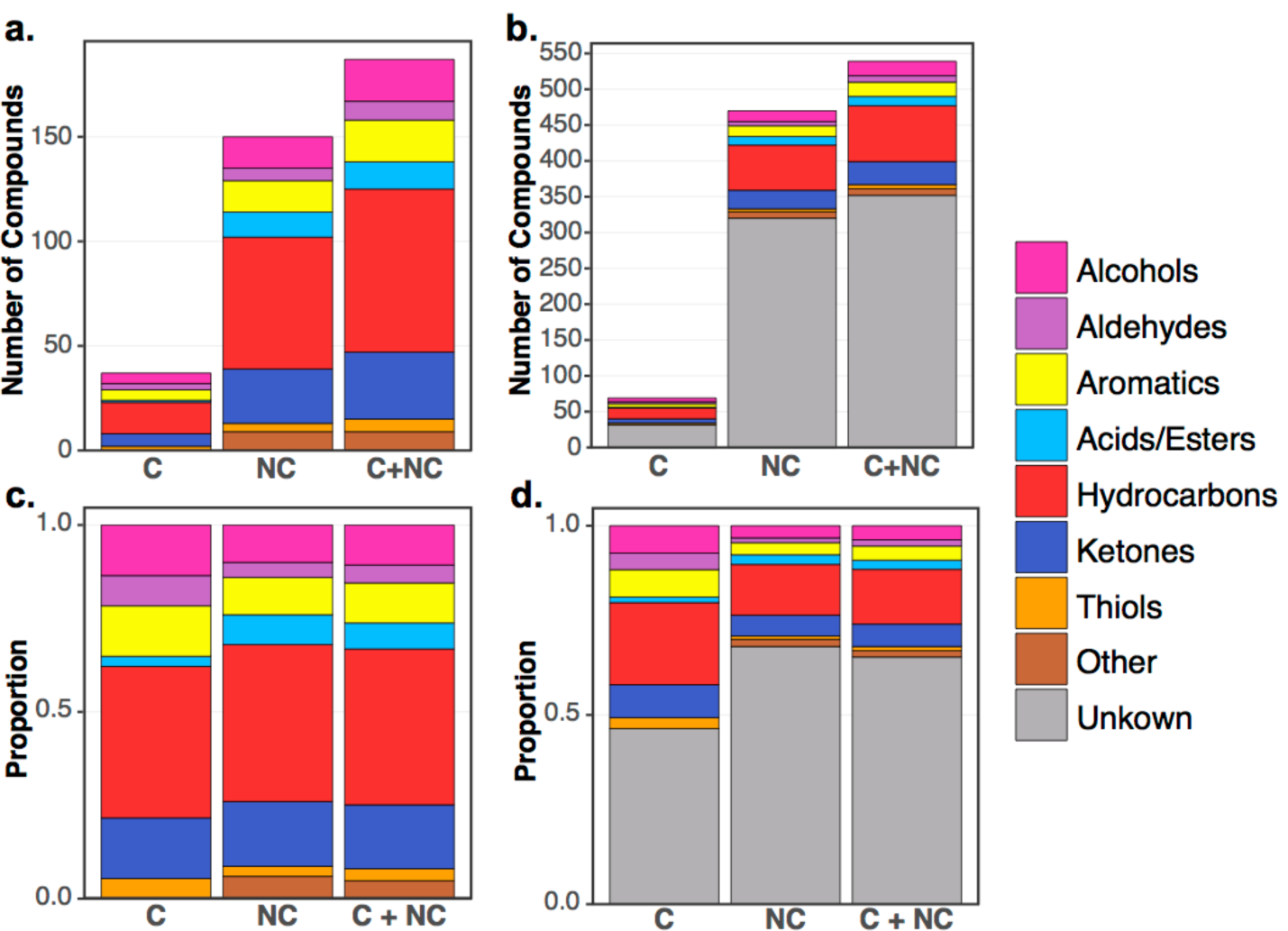
Size and chemical composition of volatile compounds metabolized by 81 *P. aeruginosa* CF chronic infection isolates. **(a)** Identified compounds; **(b)** Identified and unknown compounds; **(c)** Identified compounds, scaled to 100%; **(d)** Identified and unknown compounds, scaled to 100%. C = Core compounds, defined as detected in 95% or more of samples; NC = Non-core compounds; C+NC = sum of Core and Non-core compounds.

### Relationships in the *P. aeruginosa* CF volatilome within and between patients

Because every patient is infected by a genetically-unique strain, we hypothesized that the dissimilarities of intra-patient volatilomes would be lower than inter-patient. Additionally, due to the greater metabolic potential of early-infection isolates with intact regulatory networks, we hypothesized that the volatilomes of inter-patient Early isolates would be more dissimilar than those of inter-patient Late isolates. We calculated the pairwise Euclidean distances as a measure of dissimilarity between all Early isolate volatilomes (“inter-Early dissimilarity”), all Late isolate volatilomes (“inter-Late dissimilarity”), and between Early and Late isolate volatilomes from the same patient (“intra-patient dissimilarity”). Significant differences between means were tested using one-way ANOVA with Tukey’s HSD multiple comparison procedure. Interestingly, the inter-Early isolate dissimilarity was the lowest (mean 35.2, median 33.8) and the inter-Late isolate dissimilarity was the highest (mean 39.1, median 39.1), and these differences were significant (95% CI: 3.05 – 4.86; *P* = 0+) (Fig. 3). Similarly, mean inter-Late volatilome dissimilarity was significantly greater than that of intra-Patient volatilomes (mean 36.4, median 35.6) (95% CI: 0.41 – 4.97; *P* = .013). Even more intriguing was that the inter-Early dissimilarity was lower than even the intra-patient dissimilarity, though this difference was not significant (95% CI: −1.11 – 3.66; *P* = .515). For comparison, we also calculated the pairwise Euclidean distances of all isolates in this study (Pooled; mean 38.0, median 37.9); the mean inter-Early and mean inter-Late dissimilarities were both significantly lower than the mean dissimilarity of the Pooled isolates (95% CIs: 1.99 – 3.77, −1.65 – 0.51; *P* = 0+).

**FIG 3.**
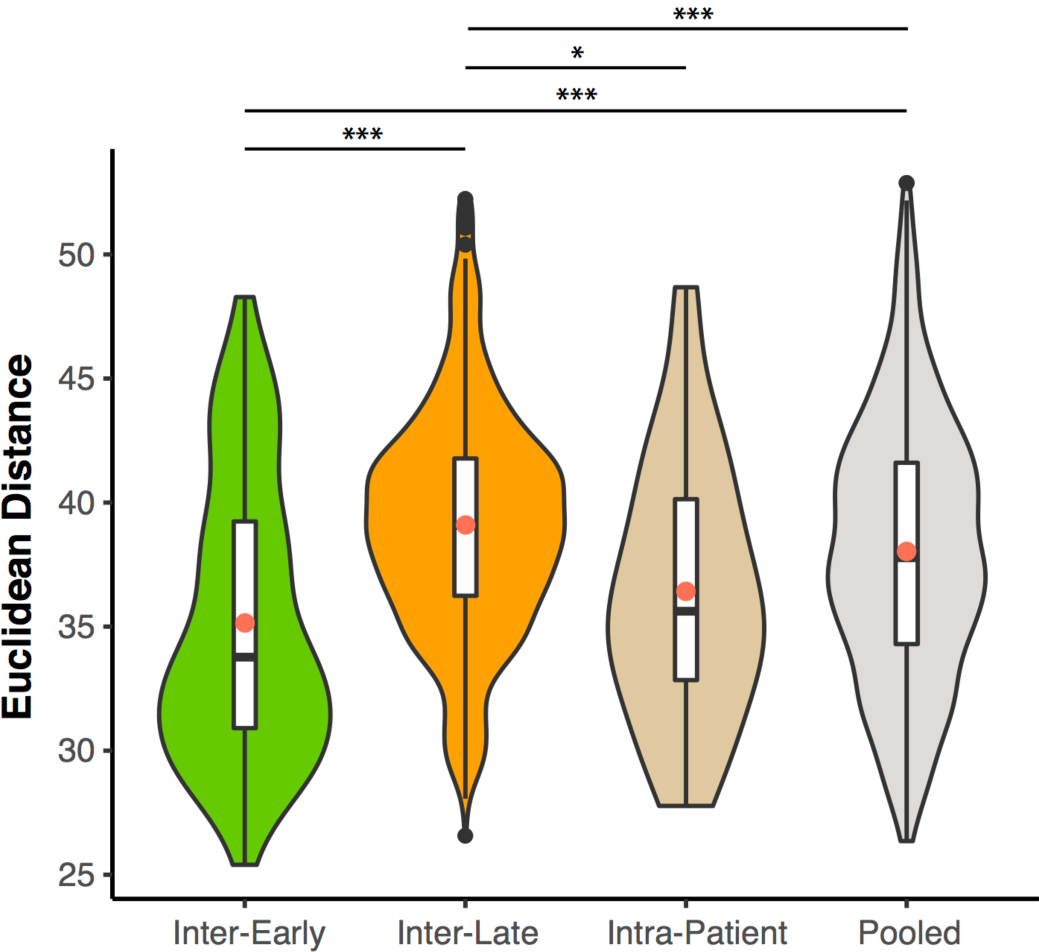
Violin plots and box plots of pairwise dissimilarities (defined as Euclidean distance) of *P. aeruginosa* chronic infection isolate volatilomes. Inter-Early are between-patient Early-Early isolate dissimilarities. Inter-Late are between-patient Late-Late isolate dissimilarities. Intra-Patient are within-patient Early-Late isolate dissimilarities. Pooled represents all pairwise volatilome dissimilarities. Red circles indicate the mean. Significant differences between means were identified using one-way ANOVA with Tukey’s HSD. \**P* < .05, \*\**P* < .01, \*\*\**P* < .001.

Visualizing the Euclidean distances between isolate volatilomes via non-metric multidimensional scaling (NMDS) and ordered dissimilarity images (ODI), we looked for clustering of clonal isolates, indicating intra-patient similarities. The NMDS and ODI of the Euclidean distances of all 81 isolate volatilomes show a primary cluster comprised mostly of intermediate isolates of Patient 23 (Fig. S1). To reduce the influence of the over-represented Patient 23 isolates in data visualization, we truncated our data set to 48 isolates by removing the Intermediate isolates of Patient 23, and the lone Patient 21 isolate. The ODI plot and NMDS of the truncated set revealed three clusters of similarity, none of which include all isolates from a single patient (Fig. 4; Fig S2a,b). This refutes our hypothesis that isolates from the same patient maintain a similar volatilome over time. To determine if any inter-patient or intra-patient isolate pairs are significantly different, we performed permutational multivariate analysis of variance (PERMANOVA) on the Euclidean distances for all possible pairs. Following Benjamini-Hochberg adjustment of *P* values, no pairs of isolates were identified as having significantly different volatilomes (*q* ≥ .1 for all isolate pairs). However, PERMANOVA conducted on all isolates indicated strong significant differences between the 81 volatilomes (*pseudo-F*_80,161_ = 28.0, *P* < .01), as well as between the volatilomes of the truncated set of 48 isolates (*pseudo-F*_47,96_ = 29.2, *P* = 0+), reinforcing the inference from the ODI and NMDS that the CF *P. aeruginosa* volatilome is heterogeneous.

**FIG 4.**
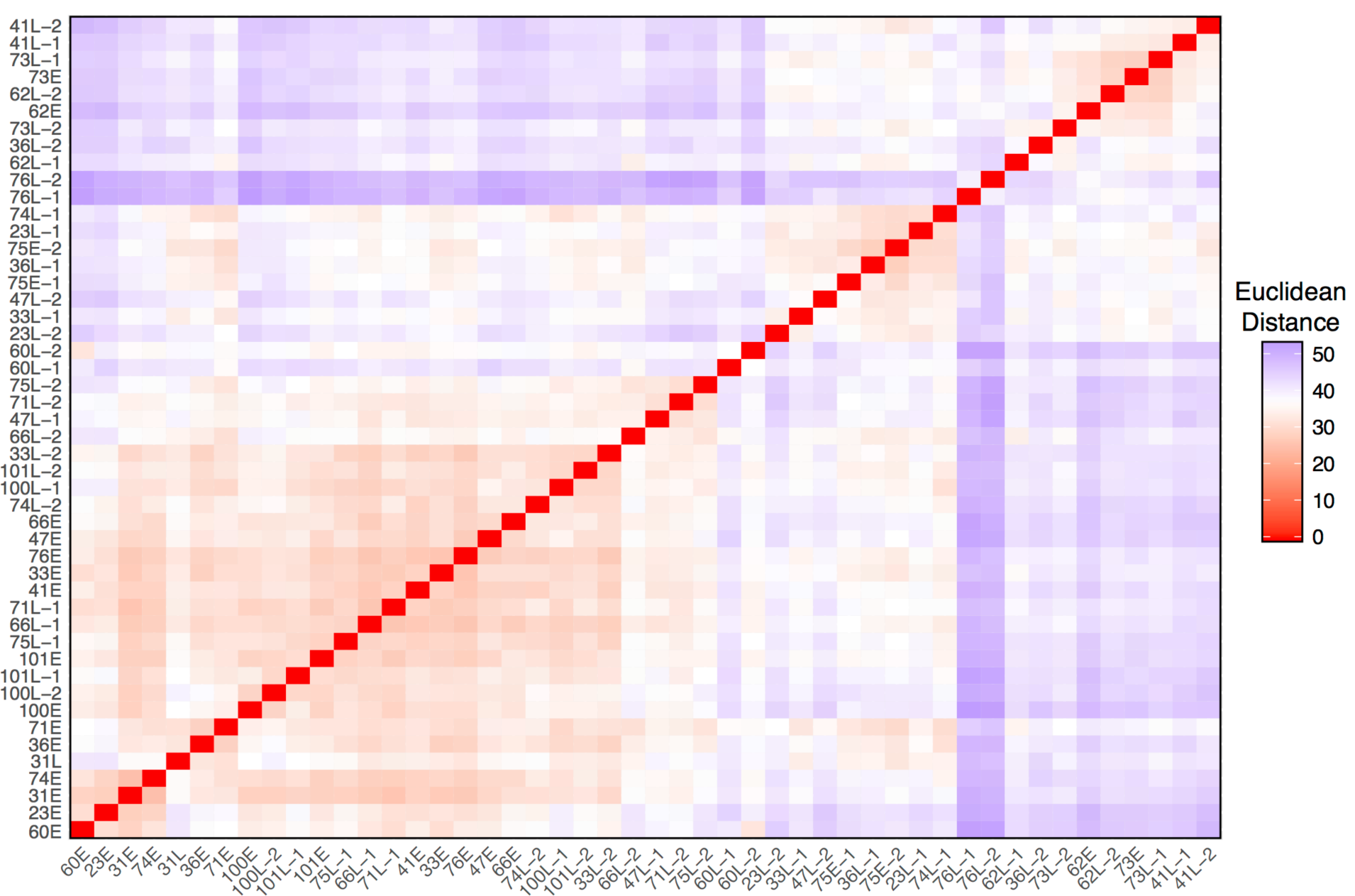
Ordered dissimilarity image (ODI) of the truncated set of 48 *P. aeruginosa* clinical isolates depicting volatilome dissimilarity defined by Euclidean distance.

Though we do not find that patient isolates maintain similar volatilomes over the duration of chronic infection, it is noteworthy that the majority of Patient 23’s isolates have strong similarities (Fig. S1). Taking a closer look at the 35 isolates from Patient 23 (Fig. 5; Fig. S2c), the ODI and NMDS revealed four distinct similarity neighborhoods: one neighborhood consisting of only the Early isolate 23E, which stands alone; a second consisting of the four latest-collected isolates 23I-32, 23I-33, 23L-1, 23L-2; a third consisting of isolates 23I-7, 23I-9, 23I-12, 23I-14, 23I-29; and a fourth encompassing the remainder of the isolates. The four latest-collected isolates are less defined by their similarity to each other than their dissimilarity to all other isolates collected during the infection. A linear regression of the Euclidean distances between the Early isolate and the Intermediate and Late isolates as a function of patient age showed that isolates became increasingly dissimilar to the Early isolate over time (Pearson’s r = 0.70, *P* = 0+) (Fig. 6). These observations are underpinned by the phenotypic differences of the four latest isolates, and the number and types of known mutations that were accumulated in the 23L versus 23E isolates (3). As described by Smith *et al*., the *P. aeruginosa* infection sampled from Patient 23 (referred to as Patient 1 in the referenced publication) diversified from patient ages 1.5 to 3 y into a population of isolates, with mutations in virulence, motility, quorum sensing, iron transport, efflux, and transcription and translation genes. The volatilome dissimilarities we measured reflect the genotypic diversification of the infection as a function of time (Fig. 6).

**FIG 5.**
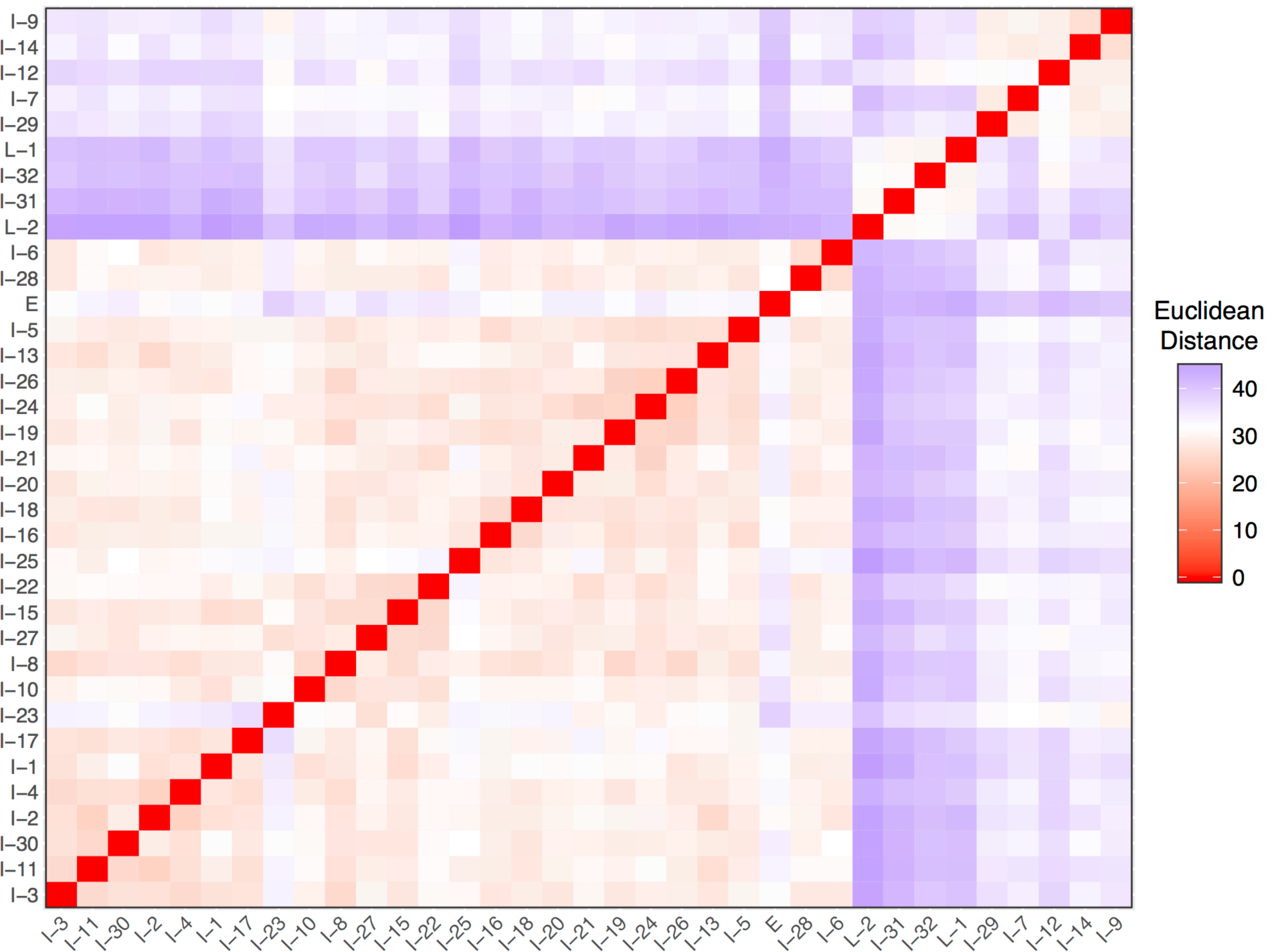
Ordered dissimilarity image (ODI) of the 35 *P. aeruginosa* clinical isolates from Patient 23 depicting volatilome dissimilarity defined by Euclidean distance. For visual clarity, the isolate labels do not include patient number (*e*.*g*., 23).

**FIG 6.**
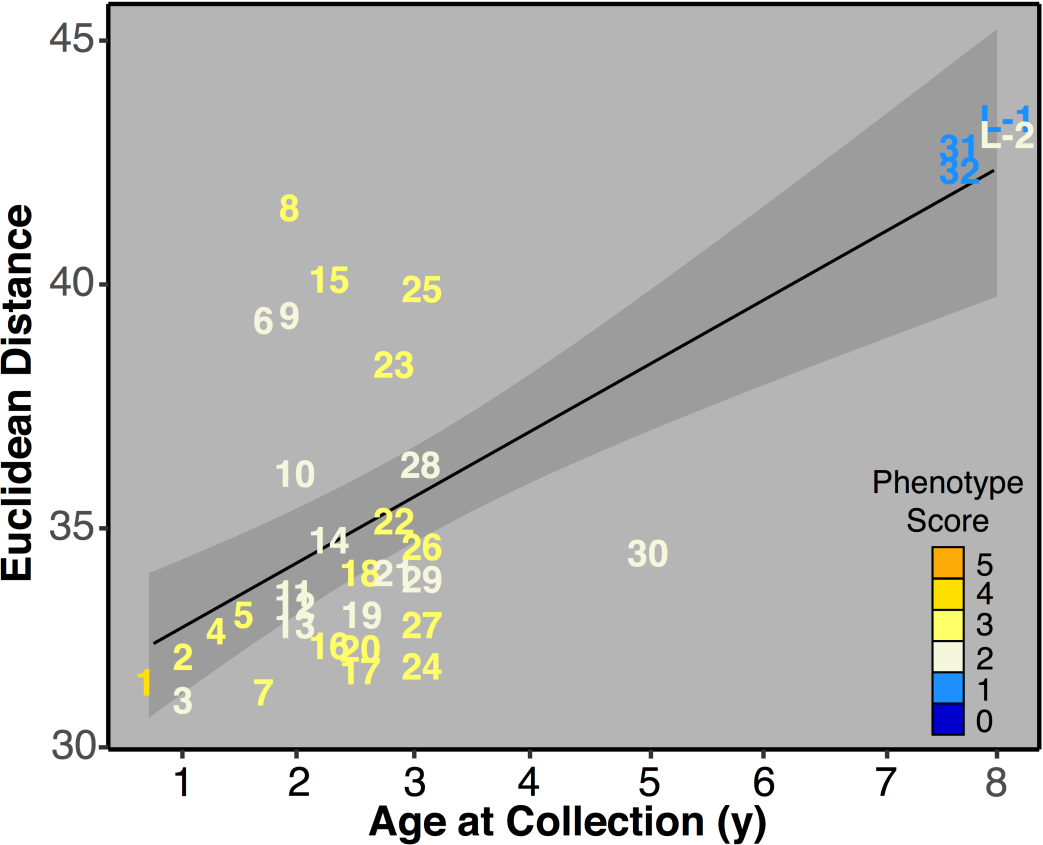
Dissimilarity of Patient 23 volatilomes over time, defined as Euclidean distance from the Early isolate (not shown). For visual clarity, the isolate labels do not include patient number (*e*.*g*., 23) and intermediate isolate labels have been truncated to only include the numeral that signifies collection order (*e*.*g*., I-1 is represented as 1). The black line represents the linear regression fit (r = 0.70, *P* = 0+), and the shaded region represents the standard error of the regression line.

### Chemical characteristics of the volatilomes of Early and Late isolates

We compared the chemical characteristics of the Early and Late volatilomes, positing that the size and chemical diversity of the volatilomes would decrease from Early to Late infection due to loss-of-function mutations. To perform the comparison, we needed to balance the size of the isolate groups, and did so by selecting ten pairs of *P. aeruginosa* isolates that had large changes in phenotypes over the duration of infection, thereby enhancing the differences we may find in the Early vs. Late volatilomes (Fig. 1; Table S1). Visual inspection of the ten pairs of GC×GC chromatograms suggests reductions in the number and variety of volatile compounds produced by Late isolates (Fig. 7, Miscellaneous Figure 2). Contrary to the appearance of the chromatograms, however, the overall size and chemical compositions of the Early and Late volatilomes were similar to each other (Fig. S3a-d), with hydrocarbons representing approximately 50% and alcohols and ketones together representing approximately 30% of the 410 Early and 441 Late volatiles. We quantified the chemical richness and diversity of the pooled Early and pooled Late isolates using the Shannon-Wiener diversity index (Table S3) and found the volatilomes of Early and Late isolates were similar, whether using all volatiles or only the non-core volatiles.

**FIG 7.**
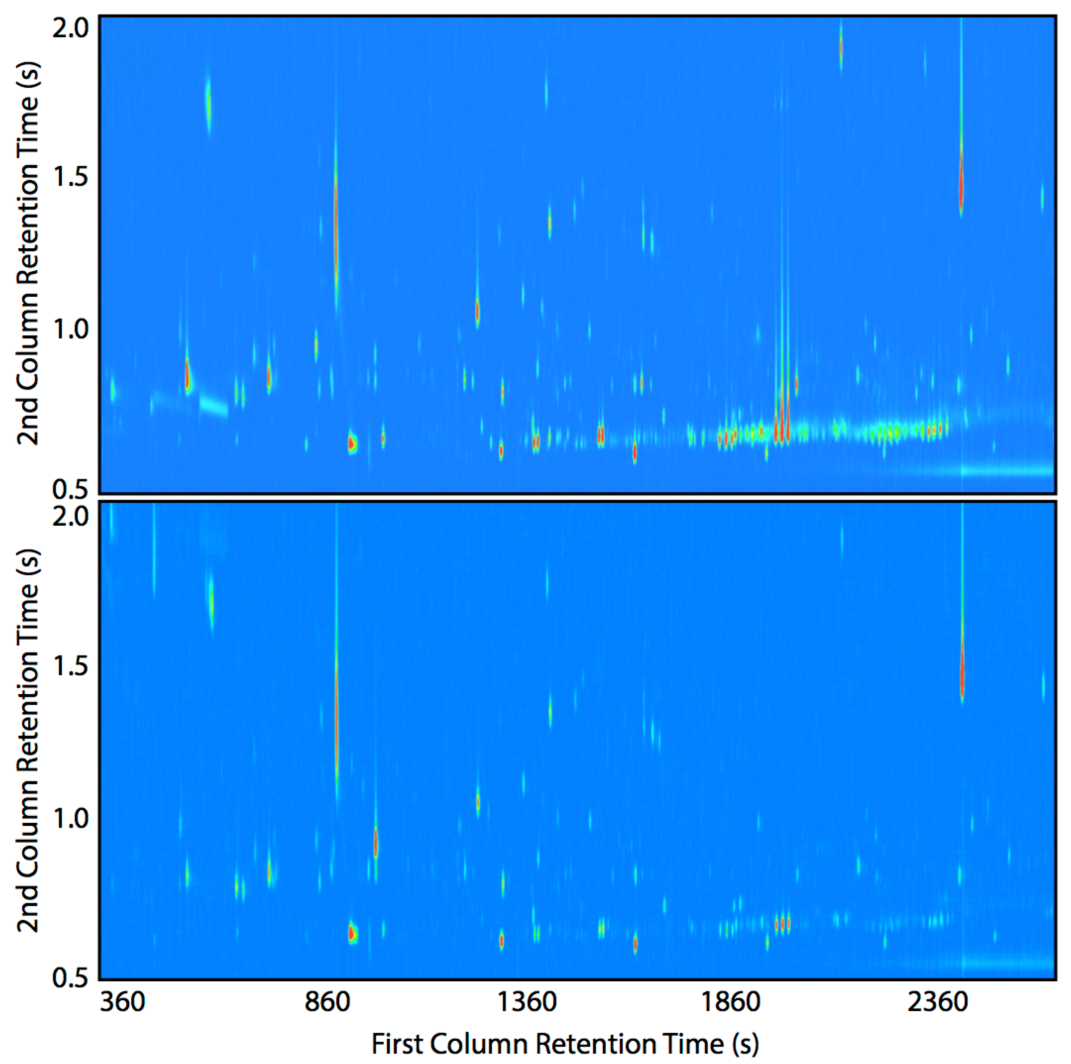
Representative GC×GC chromatograms of Early (top) and Late (bottom) chronic infection isolates (41E and 41L-2 are depicted). Dark blue represents the baseline, and peak intensity is depicted using a color gradient from light blue (low) to dark red (high). Chromatographic regions of ^1^*t*_R_ < 200 s and ^2^*t*_R_ < 0.5 s were excluded for visual clarity. All chromatograms for the ten selected Early and Late isolate pairs are provided in Supplementary Fig. 7.

The uniformity in the Early versus Late volatilomes is not a function of data aggregation, as the volatilomes of the individual isolates are highly similar in number and chemical composition, with the exception of one Late isolate (76L) that has a slow-growth phenotype (Fig. S3e-h). Furthermore, when looking at individual isolate pairs, for the vast majority there were no differences in the richness and diversity of their volatilomes (Table S3). Together, these results suggest that there are not major differences between the number and diversity of chemical compounds when comparing Early and Late infection isolates. Hierarchical clustering analysis (HCA) of the *P. aeruginosa* isolates by their volatilomes underscores this observation; we observed no discernible clustering of the Early versus Late isolates based on the presence and absence of volatiles (Fig. S4a).

### Abundance of volatile compounds in Early and Late isolates

Though we did not observe any differences in the Early and Late isolate volatilomes based on presence and absence of metabolites, the chromatograms suggest a reduced volatilome in late infection. We therefore posited that while the numbers of volatiles do not significantly change from Early to Late infection isolates, the relative concentrations of volatiles do. Using volatile abundances, we performed HCA on the 20 selected Early and Late isolates and found clustering by the time of isolate collection and phenotype score (Fig. S4b). We were interested in whether or not clustering still occurs in the larger, less curated set of 48 isolates (*i*.*e*., the truncated set), which has more discordance between phenotype and time of collection (Fig. 1). HCA of the truncated set of isolates shows that Early isolates are generally more similar to one another than Late isolates are (Fig. 8), reinforcing observations made by calculating Euclidean distances; however, significant proportions of the Early and the Late isolates are mis-clustered. Interestingly, the majority of the Late isolates clustering with the Early isolates in the upper clade have more early-like phenotypes, and several Early isolates that have more late-like phenotypes (*e*.*g*., 62E, 73E, 70E) cluster in the lower clade with Late isolates, indicating a relationship between the isolate phenomes and volatilomes. We also observed clustering by phenotype when the volatilomes of all 81 isolates were analyzed (Fig. S5). Similar phenomena are observed when clustering the truncated set or all 81 isolates by the core volatilome (Fig. S6), and even when using only the 23 core alcohols, aldehydes, and hydrocarbons (Fig. 9; Fig. S7), suggesting that a set of conserved volatiles could be identified and used as biomarkers for detecting phenotypic changes in chronic infections.

**FIG 8.**
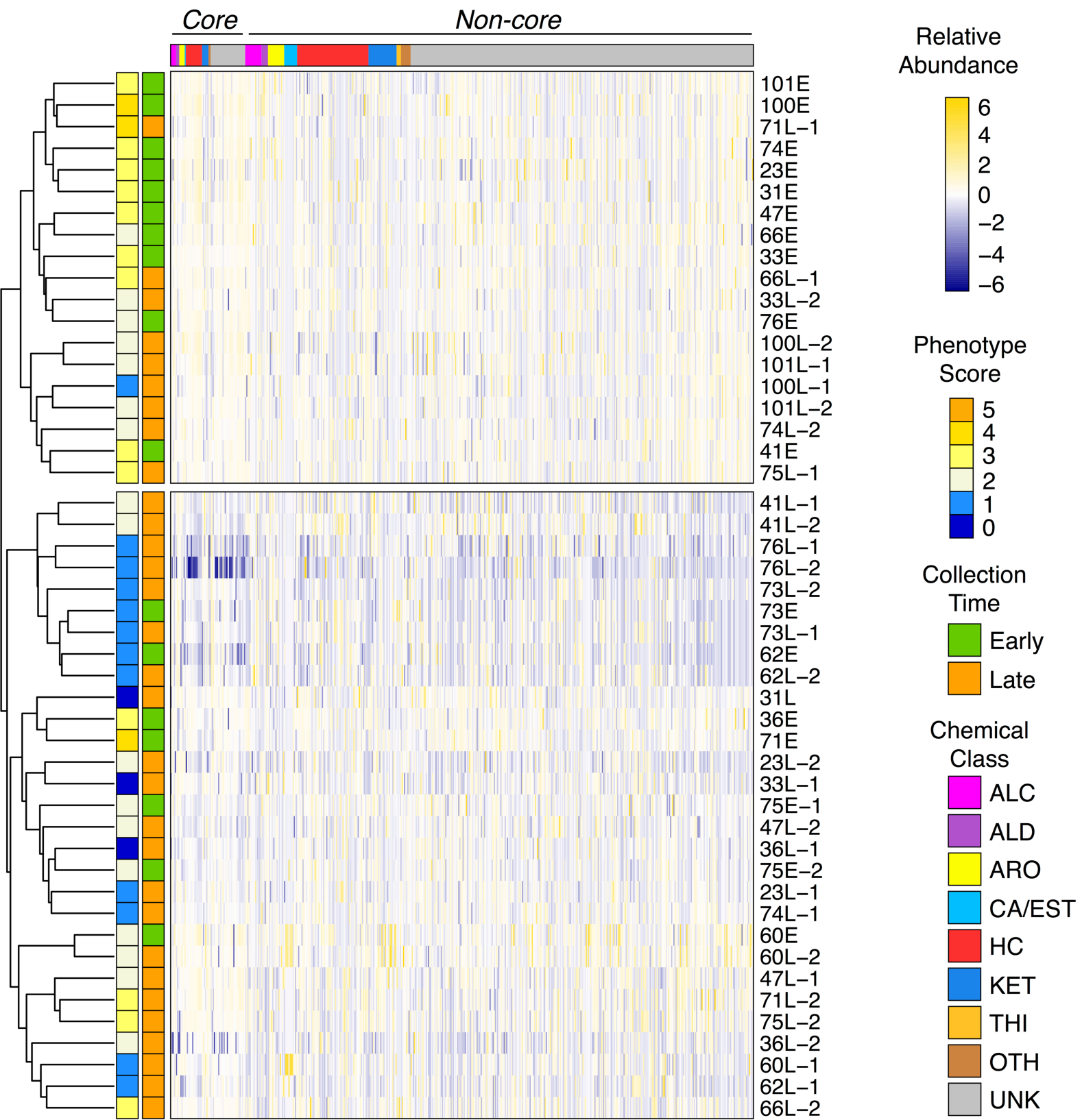
Hierarchical clustering analysis (HCA) of the truncated set of 48 *P. aeruginosa* clinical CF isolates, based on the relative abundance of 539 volatile compounds. Volatiles are in columns (standardized relative abundance). Clustering is based on rows (isolates), which are color-coded by their phenotype score (left color block) and relative time of collection (right color block). ALC = alcohols, ALD = aldehydes, ARO = aromatics, CA/EST = carboxylic acids and esters, HC = hydrocarbons, KET = ketones, THI = thiols, OTH = other compounds, UNK = unknown identity.

**FIG 9.**
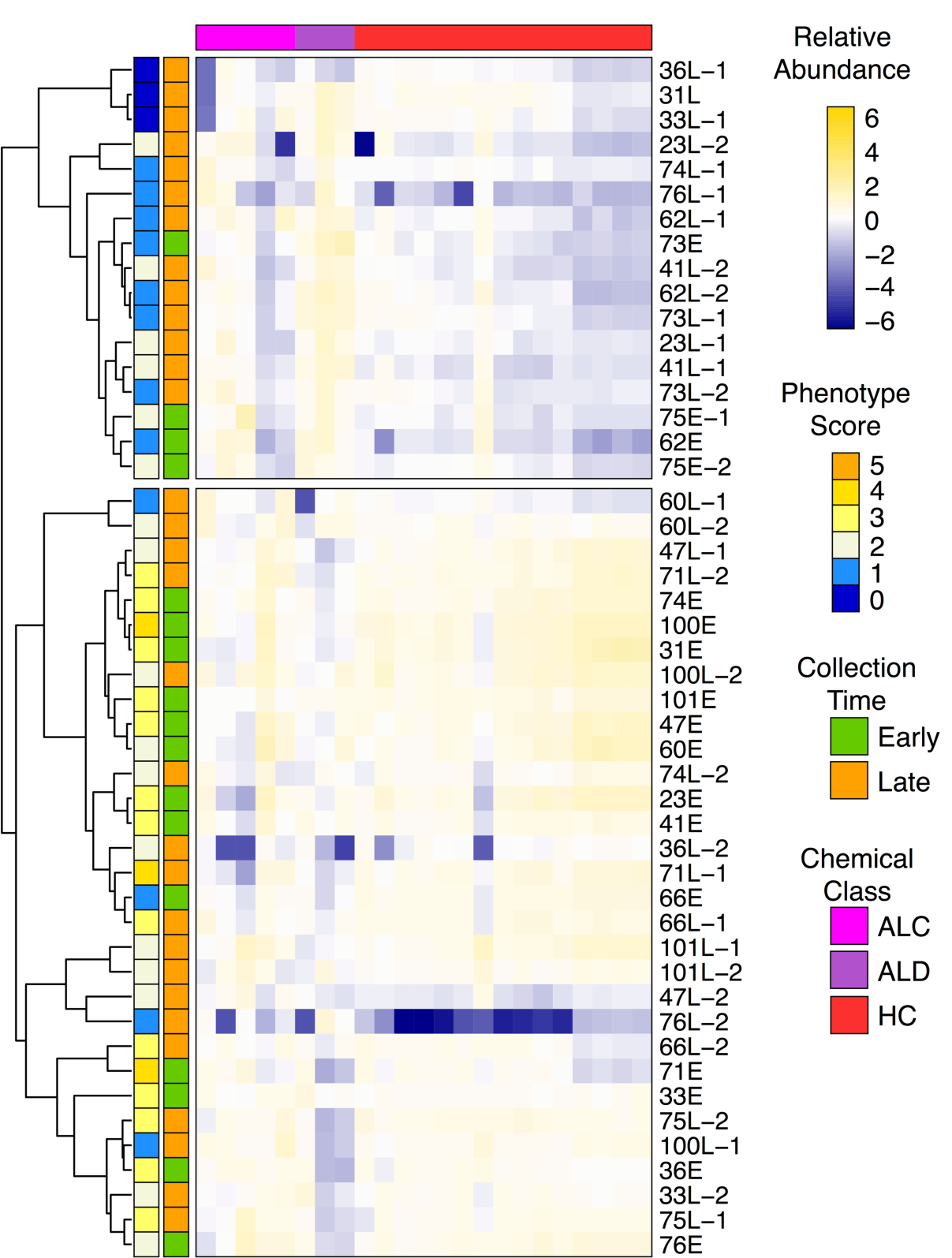
Hierarchical clustering analysis (HCA) of the truncated set of 48 *P. aeruginosa* clinical CF isolates, based on the relative abundance of 23 core alcohols, aldehydes, and hydrocarbons. Volatiles are in columns (standardized relative abundance). Clustering is based on rows (isolates), which are color-coded by their phenotype score (left color block) and relative time of collection (right color block). ALC = alcohols, ALD = aldehydes, HC = hydrocarbons.

## DISCUSSION

*P. aeruginosa* isolates from cystic fibrosis lung infections have a large and diverse volatilome. We conservatively attribute 539 detected volatiles to *P. aeruginosa* growth and metabolism, only 13% of which were core volatiles. For volatiles that we could assign to a chemical class, hydrocarbons represented the highest proportion, followed by ketones, which we observed both in the pan-volatilome and for individual patient isolates. Approximately 40% of core volatiles have been identified in other *P. aeruginosa* metabolomic studies, including the most frequently detected *in vitro* volatiles 2-butanone, 2-nonanone, dimethyl sulfide, and dimethyl disulfide (see Table 1 for references). The majority of the core volatiles we detected, however, are reported here for the first time. This opens up speculation that there may be a unique core volatile profile for strains colonizing the CF lung, though due to the inter-strain variation in the *P. aeruginosa* volatilome, many more CF and non-CF isolates will need to be characterized via untargeted GC×GC analyses to determine if this may be the case. To date, a number of studies have attempted to identify collections of volatiles to differentiate *P. aeruginosa*-positive or -negative subjects from breath (58, 74-81) and lavage (25) with promising results. Compared to this study, however, the referenced analyses were either targeted at single compounds (e.g., hydrogen cyanide or 2-aminoacetophenone), did not characterize the volatilome of *P. aeruginosa* in detail, and/or did not draw inferences to infection stage or changes across time. This study represents the first untargeted comparative analysis of *P. aeruginosa* volatilomes over long-term chronic infections, expanding the body of knowledge on *P. aeruginosa* metabolism and broadening the potential applications for breath-based diagnostics.

Despite the trend toward reduced or loss-of-phenotype expression in late infection isolates – which we hypothesized would correspond to a reduced metabolome size – we observed that Late isolates produce the same number and variety of volatile compounds as Early isolates. However, the relative abundance of volatiles produced by the Late isolates is significantly reduced. Our data suggest that mutations that arise in *P. aeruginosa* chronic infection isolates do not result in complete inhibition of metabolic pathways, but may instead reduce metabolic flux. This observation has implications for diagnostics, signifying that presence and absence of *P. aeruginosa* volatile metabolites would provide little diagnostic value for tracking chronic lung infections. Rather, a breath-based diagnostic for monitoring *P. aeruginosa* adaptations in CF will likely need to measure the relative abundance of metabolites. Importantly, a subset of core volatiles can be used to cluster isolates based on commonalities in phenotypes, indicating that a diagnostic based upon the relative abundances of conserved hydrocarbons, alcohols, and aldehydes (or a few selected volatiles from these classes) may be sufficient to track *P. aeruginosa* infection in most patients. It should be noted that core *P. aeruginosa* volatiles are not necessarily unique to *P. aeruginosa*. In fact, a review of the mVOC 2.0 microbial volatile organic compound database (82) shows nearly half of the core volatiles (and a third of named compounds overall) have been reported as being produced by other CF lung bacterial pathogens, including *Staphylococcus aureus, Klebsiella pneumoniae, Haemophilus influenzae, Stenotrophomonas maltophilia*, and *Burkholderia spp*. This further highlights the necessity of developing a breath-based diagnostic based on the relative abundances of sets of volatiles, which has been shown to accurately discriminate between human pathogens (distantly or closely related to *P. aeruginosa*) in numerous *in vitro, ex vivo*, and *in vivo* analyses (25, 62, 83-85).

Based on the genomic heterogeneity of *P. aeruginosa* (3, 33-35), we hypothesized that every patient would have a unique *P. aeruginosa* volatilome, and that intra-patient isolates would have more similar volatilomes than inter-patient isolates. To the contrary, we observed that the volatilomes of Early and Late isolates from a single patient are no more similar to one another than they are to any other isolate (intra-patient vs. pooled data, Fig. 3). Interestingly, the volatilomes of inter-patient Early isolates are even more similar than intra-patient isolates, while the volatilomes of inter-patient Late isolates have the highest dissimilarity. These data indicate that the genomic similarity within patient is less influential on the volatilome than the post-transcriptional diversity that evolves within a chronic *P. aeruginosa* infection population. The data from Patient 23, however, suggest that intra-patient isolates are more likely to have similar volatilomes when there are fewer mutational and phenotypic differences between them. From these results, we propose that a breath-based diagnostic to track *P. aeruginosa* chronic lung infections would be personalized, with each patient’s breathprint at initial infection serving as a baseline to which subsequent breathprints are compared. Sustained, significant deviations from the patient’s baseline breathprint would indicate that the *P. aeruginosa* population phenotype has changed, and therefore the patient may be at higher risk for associated poor clinical outcomes (*e*.*g*., higher risk of exacerbation) (15) and at higher risk of infection eradication failure after antibiotic treatment (12). In a longitudinal case study of a single CF patient, we have shown that changes in the dominant species colonizing the lung correlate to changes in the volatile metabolites detected in sputum (86), and therefore monitoring for breathprint deviations may also work for *P. aeruginosa* infections, as these data suggest. Despite this, the issue of specificity becomes a concern unless biomarkers for individual *P. aeruginosa* phenotypes are identified and used for tracking; this is the subject of ongoing work.

We highlight some limitations in this study. Although we make reference to Early isolates and those phenotypes associated with early infection, it is important to recognize that these isolates were merely the first culturable *P. aeruginosa* from the patients included in this study. It is likely that some patients had *P. aeruginosa* in the lung for some years in spite of persistent negative cultures into late childhood or adolescence (15), exemplified by isolates 62E and 73E exhibiting late-like phenotypes. Phenotypes discordant with collection time or apparent infection stage underscores a major drawback of culture-based methods for infection staging (*e*.*g*., *in vitro* phenotyping) in that the respiratory microbiome is prone to under-sampling, especially during initial colonization, due to the complex structure of the lung. The increased use of new therapeutics (*e*.*g*., CFTR modulators) also impacts culture-based diagnostics as they reduce sputum production, limiting the opportunity for clinical cultures. Utilizing breath-based biomarkers, on the other hand, can overcome these challenges as breath represents a sample of the entire ventilated lung environment.

*P. aeruginosa* chronic infection phenotypes are positively correlated with advanced patient age (15, 87-89), and we observe a moderate, but significant correlation (Kendall’s tau – 0.41, *P* < .001) between subject age and late-infection phenotype scores in this study. The variability of isolate volatilomes we observed might, therefore, be explained by the age of their corresponding patients. Testing this would require analyzing isolate volatilomes from a larger number of subjects that were diagnosed with their first *P. aeruginosa* infections at younger and older ages. In either case, the issue of the potential covariance between age and volatilome on *P. aeruginosa* diagnostics could be overcome by taking a multifactorial approach as opposed to volatile biomarkers only. We may find that using a combination of several clinical predictors such as age, *P. aeruginosa* phenotype, and volatilome together provides more diagnostic power than any individual metric. It will be important to explore this more closely in future work. Additionally, while we only presented volatilome trends in relation to five clinically-relevant phenotypes in this study (production of pyocyanin, rhamnolipids, and proteases, twitching mobility, and mucoidy), there are additional clinically-relevant phenotypes, including antibiotic resistance, that should be included in future analyses.

In summary, current methods for tracking *P. aeruginosa* lung infection progression are predominantly culture-based (16, 21), and our results suggest a potential role for *in vitro* volatile metabolomics in staging chronic infections in conjunction with established clinical laboratory methods. The clinically-correlated *P. aeruginosa* phenotypes we measured in this study – mucoidy, motility, and quorum-regulated traits – are sensitive to environment (90-93); as such, the metabolomes we characterized from *in vitro* cultures are unlikely to directly reflect the metabolomes these isolates produced in the CF lung. For the development of a breath test for tracking infections *in situ*, it will be necessary to utilize culture models that are more reflective of the lung (94-97) and to collect *ex vivo* and *in vivo* infection volatilomes (25, 75, 98). The utility of breath for tracking chronic lung infections is being demonstrated in tuberculosis lung infection models (26), and therefore we anticipate that the development of a breath-based diagnostic for tracking *P. aeruginosa* chronic lung infections is feasible. It will also be important to ensure that the *P. aeruginosa* volatiles we have identified are detectable in breath. This is the primary goal of the observational clinical study IMproving *P. Aeruginosa* deteCTion using Breath (IMPACT-Breath) that is underway. Despite the limitations of *in vitro* analyses, this study presents the first comprehensive analysis of the *P. aeruginosa* volatilome from chronic CF lung infections, and lays the groundwork for the application of volatile metabolomics in tracking CF lung disease.

## Supporting information

Miscellaneous Information

## ACKNOWLEDGEMENTS

Funding to support this work was provided by the Cystic Fibrosis Foundation (Hill17P0, Hill18A0-CI), the National Institutes of Health (R56HL139846), and the ASU School of Life Sciences Undergraduate Research (SOLUR) program. Isolates were obtained from the Cystic Fibrosis Isolate Core at Seattle Children’s Research Institute, funded by the National Institutes of Health (P30 DK089507) and the Cystic Fibrosis Foundation (HOFFMA20Y2-OUT).

## AUTHOR CONTRIBUTIONS

HB conceived of the study; HB, TD designed the experiments; AK, CB, SW, and TD collected phenotype data; TD collected metabolomics data; TD and MR processed and analyzed the metabolomics data; HB and TD wrote and edited the manuscript; all authors approved the final version.

---

**FIG S1**. Non-metric multidimensional scaling (NMDS) plots of 81 *P. aeruginosa* clinical isolates depicting volatilome dissimilarity defined by Euclidean distance **(a)** colored by collection time and **(b)** colored by patient; **(c)** Ordered dissimilarity image (ODI) of 81 *P. aeruginosa* clinical isolates depicting volatilome dissimilarity defined by Euclidean distance.

**FIG S2**. Non-metric multidimensional scaling (NMDS) plots of the truncated set of 48 *P. aeruginosa* clinical isolates depicting volatilome dissimilarity defined by Euclidean distance **(a)** colored by collection time and **(b)** colored by patient; **(c)** NMDS plot of the 35 *P. aeruginosa* clinical isolate volatilomes from Patient 23 depicting volatilome dissimilarity defined by Euclidean distance.

**FIG S3**. Size and chemical composition of volatile compounds metabolized by the selected ten Early-Late pairs of *P. aeruginosa* chronic infection CF isolates. **(a)** Identified compounds; **(b)** Identified and unknown compounds; **(c)** Identified compounds, scaled to 100%; **(d)** Identified and unknown compounds, scaled to 100%. Size and chemical composition of volatile compounds metabolized by each of the ten Early-Late pairs of *P. aeruginosa* chronic infection CF isolates. **(e)** Identified compounds; **(f)** Identified and unknown compounds; **(g)** Identified compounds, scaled to 100%; **(h)** Identified and unknown compounds, scaled to 100% C = Core compounds, defined as detected in 95% or more of samples; NC = Non-core compounds; C+NC = sum of Core and Non-core compounds.

**FIG S4**. Hierarchical clustering analysis (HCA) of the ten Early-Late pairs of *P. aeruginosa* clinical CF isolates, based on **(a)** presence and absence and **(b)** relative abundance of 472 volatile compounds. Core compounds are present in at least 95% of isolates; Non-core compounds are present in less than 95%. Volatiles are in columns. Clustering is based on rows (isolates), which are color-coded by their phenotype score (left color block) and relative time of collection (right color block).

**FIG S5**. Hierarchical clustering analysis (HCA) of the 81 *P. aeruginosa* clinical CF isolates, based on the relative abundance of 539 volatile compounds. Core compounds are present in at least 95% of isolates; Non-core compounds are present in less than 95%. Volatiles are in columns (standardized relative abundance). Clustering is based on rows (isolates), which are color-coded by their phenotype score (left color block) and relative time of collection (right color block).

**FIG S6**. Hierarchical clustering analysis (HCA) of **(a)** the truncated set of 48 and **(b)** full set of 81 *P. aeruginosa* clinical CF isolates, based on the relative abundance of the 69 core volatile compounds. Volatiles are in columns (standardized relative abundance). Clustering is based on rows (isolates), which are color-coded by their phenotype score (left color block) and relative time of collection (right color block).

**FIG S7**. Hierarchical clustering analysis (HCA) of the 81 *P. aeruginosa* clinical CF isolates, based on the relative abundance of 23 core alcohols, aldehydes, and hydrocarbons. Volatiles are in columns (standardized relative abundance). Clustering is based on rows (isolates), which are color-coded by their phenotype score (left color block) and relative time of collection (right color block).

