## Supplementary Information for "*PSEUDOMONAS AERUGINOSA* VOLATILOME CHARACTERISTICS AND ADAPTATIONS IN CHRONIC CYSTIC FIBROSIS LUNG INFECTIONS"

**Supplementary Table 1.** Summary statistics of 81 *P. aeruginosa* isolates collected from 17 patients with CF. Isolate 21 was the only isolate from that patient included in this study. E = isolate collected early in chronic infection; L = isolate collected late in chronic infection; I = isolate collected intermediately between the early and late isolates. AGE = age in years of patient when isolate was collected. PYO = pyocyanin (absorbance); PRO = protease (cm); RHL = rhamnolipids (score); TWI = twitching motility (cm); MUC = mucoidy (visual score). See PART A of this document for information on the assays used to collect phenotype data, which are presented in the table as raw, non-scaled values that are the mean of three biological replicates. "Score" was determined, for each replicate, by scaling the data for each phenotype to a range of 0 to 1, where 1 is the maximum value and 0 is the minimum value (excluding outliers). Scaled phenotype data were then summed to yield the Score. Age is not included in calculation of Scores. The Score reported in the table is the mean of triplicate Scores from the three biological replicates, rounded to the nearest integer. *P* values indicate the significance of the late isolate Score when compared to the early isolate Score using the Wilcoxon signed-rank test (with continuity correction where appropriate). Isolates selected for further Early vs. Late analyses are indicated with asterisks (*).

| Isolate | Isolate Core Code[a] | E/L/I | AGE | PYO | PRO | RHL | TWI | MUC | Score | Significance |
|---|---|---|---|---|---|---|---|---|---|---|
| **21** | | N/A | N/A | 0.049 | 0.00 | 1 | 0.05 | 0.5 | 1 | |
| **23E*** | AMT0023-30 | E | 0.5 | 0.609 | 1.08 | 5 | 0.43 | 0.92 | 3 | |
| **23I-1** | AMT0023-10 | I | 0.75 | 0.661 | 1.23 | 4 | 0.54 | 0.83 | 4 | *P* = 1 |
| **23I-2** | AMT0023-33 | I | 1 | 0.592 | 0.77 | 4 | 0.07 | 1.00 | 3 | *P* ≈ .37 |
| **23I-3** | AMT0023-25 | I | 1 | 0.576 | 0.83 | 3 | 0.10 | 1.00 | 2 | *P* ≈ .35 |
| **23I-4** | AMT0023-26 | I | 1.25 | 0.793 | 0.85 | 3 | 0.00 | 1.00 | 3 | *P* ≈ .15 |
| **23I-5** | AMT0023-12 | I | 1.5 | 0.754 | 1.17 | 3 | 0.00 | 1.00 | 3 | *P* = .59 |
| **23I-6** | AMT0023-20 | I | 1.75 | 0.867 | 0.43 | 3 | 0.03 | 1.00 | 2 | *P* = .15 |
| **23I-7** | AMT0023-18 | I | 1.75 | 0.566 | 0.72 | 4 | 0.10 | 1.00 | 3 | *P* ≈ .15 |
| **23I-8** | AMT0023-2 | I | 2 | 0.700 | 1.10 | 4 | 0.00 | 1.00 | 3 | *P* ≈ 1 |
| **23I-9** | AMT0023-27 | I | 2 | 0.491 | 0.53 | 3 | 0.00 | 1.00 | 2 | *P* ≈ .17 |
| **23I-10** | AMT0023-14 | I | 2 | 0.531 | 1.35 | 3 | 0.00 | 0.92 | 4 | *P* ≈ .37 |
| **23I-11** | AMT0023-21 | I | 2 | 0.530 | 0.87 | 2 | 0.03 | 1.00 | 3 | *P* ≈ .17 |
| **23I-12** | AMT023-11 | I | 2 | 0.565 | 0.37 | 3 | 0.00 | 1.00 | 2 | *P* = .17 |
| **23I-13** | AMT0023-22 | I | 2 | 0.622 | 0.60 | 3 | 0.00 | 1.00 | 2 | *P* ≈ .15 |
| **23I-14** | AMT0023-23 | I | 2.25 | 0.457 | 0.00 | 3 | 0.00 | 1.00 | 2 | *P* ≈ .17 |
| **23I-15** | AMT0023-19 | I | 2.25 | 0.862 | 1.20 | 4 | 0.00 | 1.00 | 3 | *P* ≈ .35 |
| **23I-16** | AMT0023-6 | I | 2.25 | 0.540 | 1.10 | 4 | 0.22 | 0.83 | 3 | *P* ≈ .77 |
| **23I-17** | AMT0023-24 | I | 2.5 | 0.586 | 0.87 | 2 | 0.48 | 0.50 | 3 | *P* ≈ .15 |
| **23I-18** | AMT0023-15 | I | 2.5 | 0.882 | 1.67 | 2 | 0.22 | 0.92 | 4 | *P* ≈ .35 |
| **23I-19** | AMT0023-9 | I | 2.5 | 0.397 | 0.95 | 3 | 0.00 | 1.00 | 2 | *P* ≈ .37 |
| **23I-20** | AMT0023-16 | I | 2.5 | 0.998 | 1.10 | 2 | 0.00 | 1.00 | 3 | *P* ≈ .35 |
| **23I-21** | AMT0023-13 | I | 2.75 | 0.837 | 1.13 | 3 | 0.00 | 1.00 | 3 | *P* ≈ .35 |
| **23I-22** | AMT0023-17 | I | 2.75 | 1.061 | 1.10 | 3 | 0.05 | 1.00 | 3 | *P* ≈ .15 |
| **23I-23** | AMT0023-7 | I | 2.75 | 0.596 | 1.20 | 4 | 0.41 | 0.92 | 3 | *P* ≈ 1 |
| **23I-24** | AMT0023-3 | I | 3 | 0.774 | 1.08 | 3 | 0.00 | 1.00 | 3 | *P* ≈ 1 |

| | | | | | | | | | | |
|---|---|---|---|---|---|---|---|---|---|---|
| **23I-25** | AMT0023-8 | I | 3 | 0.676 | 1.30 | 3 | 0.39 | 0.92 | 3 | *P* ≈ 1 |
| **23I-26** | AMT0023-4 | I | 3 | 0.530 | 1.02 | 3 | 0.48 | 1.00 | 3 | *P* = 1 |
| **23I-27** | AMT0023-5 | I | 3 | 0.527 | 1.02 | 3 | 0.03 | 1.00 | 3 | *P* = .35 |
| **23I-28** | AMT0023-1 | I | 3 | 0.538 | 0.93 | 3 | 0.00 | 1.00 | 3 | *P* ≈ .37 |
| **23I-29** | AMT0023-28 | I | 3 | 0.785 | 0.27 | 4 | 0.17 | 1.00 | 2 | *P* = .37 |
| **23I-30** | AMT0023-32 | I | 5 | 0.711 | 0.80 | 2 | 0.05 | 0.83 | 2 | *P* = .37 |
| **23I-31** | AMT0023-31 | I | 7.6 | 0.030 | 0.30 | 1 | 0.00 | 1.00 | 1 | *P* ≈ .25 |
| **23I-32** | AMT0023-29 | I | 7.6 | 0.066 | 0.32 | 1 | 0.00 | 0.92 | 1 | *P* ≈ .25 |
| **23L-1*** | AMT0023-34 | L | 8 | 0.042 | 0.22 | 2 | 0.05 | 0.92 | 1 | *P* ≈ .17 |
| **23L-2** | AMT0023-35 | L | 8 | 0.174 | 0.30 | 2 | 0.00 | 1.00 | 2 | *P* = .25 |
| **31E*** | AMT0031-2 | E | 6.3 | 0.840 | 0.93 | 5 | 0.21 | 0.75 | 3 | |
| **31L-1*** | AMT0031-1 | L | 12.8 | 0.106 | 0.00 | 0 | 0.17 | 0.00 | 0 | *P* ≈ .25 |
| **33E*** | AMT0033-2 | E | 1.1 | 0.614 | 0.77 | 5 | 0.38 | 0.75 | 3 | |
| **33L-1** | AMT0033-1 | L | 13.2 | 0.003 | 0.00 | 0 | 0.00 | 0.17 | 0 | *P* = .25 |
| **33L-2*** | AMT0033-3 | L | 13.2 | 0.145 | 0.23 | 1 | 0.20 | 1.00 | 2 | *P* = .37 |
| **36E*** | AMT0036-3 | E | 6.7 | 0.902 | 0.65 | 5 | 0.00 | 1.00 | 3 | |
| **36L-1*** | AMT0036-1 | L | 15.4 | 0.531 | 0.00 | 0 | 0.00 | 0.25 | 0 | *P* = .17 |
| **36L-2** | AMT0036-2 | L | 15.4 | 1.492 | 0.00 | 2 | 0.00 | 1.00 | 2 | *P* = .15 |
| **41E*** | AMT0041-1 | E | 5.6 | 0.636 | 0.87 | 4 | 0.03 | 1.00 | 3 | |
| **41L-1** | AMT0041-2 | L | 12.8 | 1.112 | 0.00 | 4 | 0.03 | 1.00 | 2 | *P* ≈ 1 |
| **41L-2*** | AMT0041-3 | L | 12.8 | 0.103 | 0.00 | 3 | 0.03 | 1.00 | 2 | *P* ≈ .37 |
| **47E** | AMT0047-2 | E | 0.8 | 0.504 | 0.57 | 5 | 0.23 | 1.00 | 3 | |
| **47L-1** | AMT0047-1 | L | 6.8 | 0.725 | 0.32 | 5 | 0.03 | 0.17 | 2 | *P* ≈ .17 |
| **47L-2** | AMT0047-3 | L | 7.3 | 0.212 | 0.00 | 5 | 0.25 | 0.75 | 2 | *P* ≈ .17 |
| **60E** | AMT0060-3 | E | 7.7 | 0.541 | 0.80 | 1 | 0.20 | 1.00 | 2 | |
| **60L-1** | AMT0060-2 | L | 15.4 | 0.267 | 0.00 | 1 | 0.10 | 0.33 | 1 | *P* = .17 |
| **60L-2** | AMT0060-1 | L | 15.4 | 0.342 | 0.45 | 2 | 0.23 | 1.00 | 2 | *P* = 1 |
| **62E** | AMT0062-2 | E | 4.4 | 0.001 | 0.00 | 0 | 0.00 | 0.92 | 1 | |
| **62L-1** | AMT0062-3 | L | 13.1 | 0.096 | 0.00 | 0 | 0.00 | 0.92 | 1 | N/A |
| **62L-2** | AMT0062-1 | L | 13.1 | 0.083 | 0.00 | 1 | 0.00 | 0.92 | 1 | N/A |
| **66E*** | AMT0066-3 | E | 7.2 | 0.569 | 0.55 | 2 | 0.20 | 1.00 | 2 | |
| **66L-1*** | AMT0066-2 | L | 13.7 | 1.233 | 0.42 | 3 | 0.23 | 0.83 | 3 | *P* ≈ 1 |
| **66L-2** | AMT0066-1 | L | 15.2 | 1.469 | 0.47 | 3 | 0.28 | 0.67 | 3 | *P* ≈ 1 |
| **71E** | AMT0071-2 | E | 3 | 1.827 | 1.70 | 4 | 0.30 | 0.92 | 4 | |
| **71L-1** | AMT0071-1 | L | 9.1 | 3.177 | 1.47 | 3 | 0.18 | 1.00 | 4 | *P* = 1 |

| | | | | | | | | | | |
|---|---|---|---|---|---|---|---|---|---|---|
| **71L-2** | AMT0071-3 | L | 9.1 | 1.611 | 1.62 | 3 | 0.37 | 0.75 | 3 | *P* ≈ 1 |
| **73E** | AMT0073-3 | E | 8.6 | 0.083 | 0.22 | 2 | 0.00 | 1.00 | 1 | |
| **73L-1** | AMT0073-2 | L | 21.1 | 0.418 | 0.00 | 1 | 0.03 | 1.00 | 1 | N/A |
| **73L-2** | AMT0073-1 | L | 21.1 | 0.041 | 0.00 | 1 | 0.03 | 1.00 | 1 | *P* ≈ .35 |
| **74E** | AMT0074-1 | E | 9.2 | 0.656 | 1.20 | 2 | 0.38 | 0.67 | 3 | |
| **74L-1** | AMT0074-2 | L | 19.6 | 0.500 | 0.00 | 1 | 0.03 | 0.50 | 1 | p ≈ .15 |
| **74L-2** | AMT0074-3 | L | 19.6 | 0.677 | 0.47 | 2 | 0.05 | 1.00 | 2 | p ≈ .35 |
| **75E-1*** | AMT0075-3 | E | 7 | 0.249 | 0.35 | 4 | 0.18 | 1.00 | 2 | |
| **75E-2** | AMT0075-1 | E | 7.1 | 0.053 | 0.20 | 4 | 0.08 | 1.00 | 2 | |
| **75L-1*** | AMT0075-2 | L | 17.6 | 0.830 | 0.58 | 4 | 0.27 | 0.83 | 3 | *P* = 1;<br>*P* = 1 |
| **75L-2** | AMT0075-4 | L | 23.4 | 0.721 | 0.38 | 4 | 0.28 | 1.00 | 3 | *P* = 1;<br>*P* = 1 |
| **76E*** | AMT0076-3 | E | 10.8 | 0.898 | 0.98 | 5 | 0.10 | 0.50 | 2 | |
| **76L-1** | AMT0076-1 | L | 19.6 | 0.108 | 0.00 | 0 | 0.00 | 1.00 | 1 | *P* = .35 |
| **76L-2*** | AMT0076-2 | L | 19.6 | 0.075 | 0.13 | 1 | 0.00 | 1.00 | 1 | *P* ≈ .37 |
| **100E*** | NC-AMT0100-2 | E | 2.3 | 0.769 | 0.68 | 5 | 0.68 | 1.00 | 4 | |
| **100L-1*** | NC-AMT0100-1 | L | 9.6 | 0.870 | 0.42 | 1 | 0.00 | 0.33 | 1 | *P* = .25 |
| **100L-2** | NC-AMT0100-3 | L | 9.6 | 0.433 | 0.50 | 5 | 0.10 | 0.92 | 2 | *P* ≈ .17 |
| **101E*** | NC-AMT0101-3 | E | 1 | 0.594 | 0.85 | 5 | 0.53 | 0.83 | 3 | |
| **101L-1** | NC-AMT0101-2 | L | 9.6 | 0.673 | 0.60 | 4 | 0.03 | 1.00 | 2 | *P* ≈ .78 |
| **101L-2*** | NC-AMT0101-3 | L | 9.6 | 0.538 | 0.03 | 3 | 0.00 | 1.00 | 2 | *P* ≈ .37 |

[a] Isolate Core Code refers to the strain name assigned by the Cystic Fibrosis Isolate Core at Seattle Children's Center for Global Infectious Disease Research. Isolates are available to researchers by request to the CF Isolate Core: https://www.seattlechildrens.org/research/resources/cystic-fibrosis-isolate/

**Supplementary Table 2.** See **Supplementary File**. Table of 539 peaks metabolized by 81 *P. aeruginosa* clinical CF isolates.

**Supplementary Table 3.** Measures of richness and Shannon diversity of the selected ten Early-Late pairs of *P. aeruginosa* clinical CF isolate volatilomes.

| Isolate | | C + NC | | Core (C) | | Non-core (NC) | |
|---|---|---|---|---|---|---|---|
| | | *Richness* | *Diversity* | *Richness* | *Diversity* | *Richness* | *Diversity* |
| **Pooled** | *Early* | 9 | 1.68 | 7 | 1.66 | 9 | 1.65 |
| | *Late* | 9 | 1.64 | 7 | 1.66 | 9 | 1.60 |
| **P23** | *Early* | 9 | 1.77 | 7 | 1.66 | 9 | 1.79 |
| | *Late* | 8 | 1.63 | 7 | 1.66 | 8 | 1.53 |
| **P31** | *Early* | 9 | 1.71 | 7 | 1.66 | 9 | 1.69 |
| | *Late* | 8 | 1.61 | 7 | 1.66 | 7 | 1.45 |
| **P33** | *Early* | 9 | 1.70 | 7 | 1.66 | 9 | 1.67 |
| | *Late* | 9 | 1.68 | 7 | 1.66 | 9 | 1.61 |
| **P36** | *Early* | 9 | 1.62 | 7 | 1.66 | 8 | 1.49 |
| | *Late* | 9 | 1.68 | 7 | 1.66 | 8 | 1.58 |
| **P41** | *Early* | 9 | 1.71 | 7 | 1.66 | 9 | 1.69 |
| | *Late* | 9 | 1.63 | 7 | 1.66 | 8 | 1.53 |
| **P66** | *Early* | 9 | 1.72 | 7 | 1.66 | 8 | 1.67 |
| | *Late* | 9 | 1.65 | 7 | 1.66 | 7 | 1.54 |
| **P75** | *Early* | 9 | 1.68 | 7 | 1.66 | 8 | 1.59 |
| | *Late* | 8 | 1.68 | 7 | 1.66 | 9 | 1.64 |
| **P76** | *Early* | 9 | 1.69 | 7 | 1.66 | 9 | 1.64 |
| | *Late* | 9 | 1.90 | 7 | 1.78 | 8 | 1.87 |
| **P100** | *Early* | 9 | 1.78 | 7 | 1.66 | 9 | 1.80 |
| | *Late* | 9 | 1.69 | 7 | 1.66 | 8 | 1.61 |
| **P101** | *Early* | 9 | 1.78 | 7 | 1.66 | 9 | 1.81 |
| | *Late* | 9 | 1.74 | 7 | 1.66 | 9 | 1.72 |

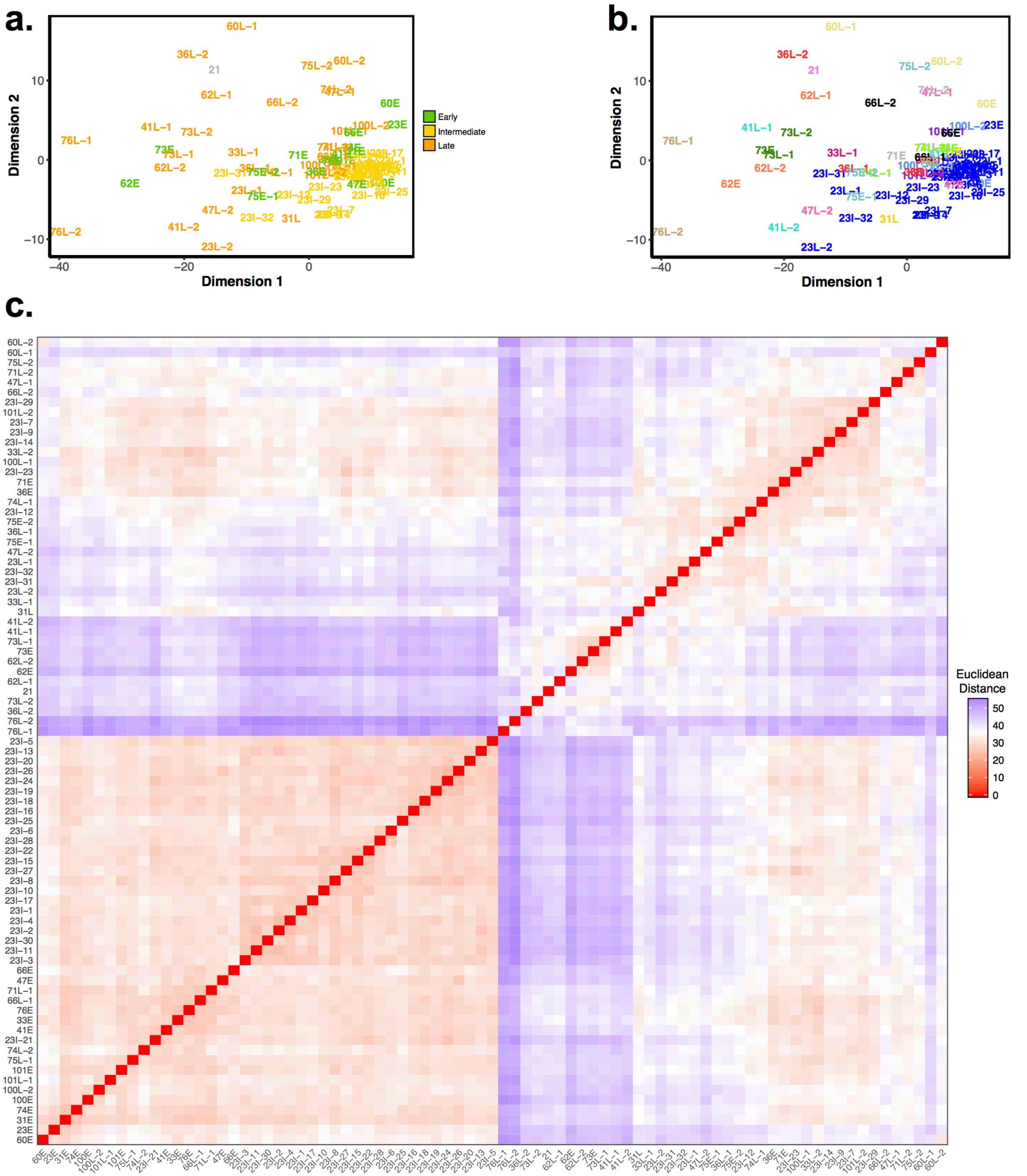

**Figure S1**. Non-metric multidimensional scaling (NMDS) plots of 81 *P. aeruginosa* clinical isolates depicting volatilome dissimilarity defined by Euclidean distance **(a)** colored by collection time and **(b)** colored by patient; **(c)** Ordered dissimilarity image (ODI) of 81 *P. aeruginosa* clinical isolates depicting volatilome dissimilarity defined by Euclidean distance.

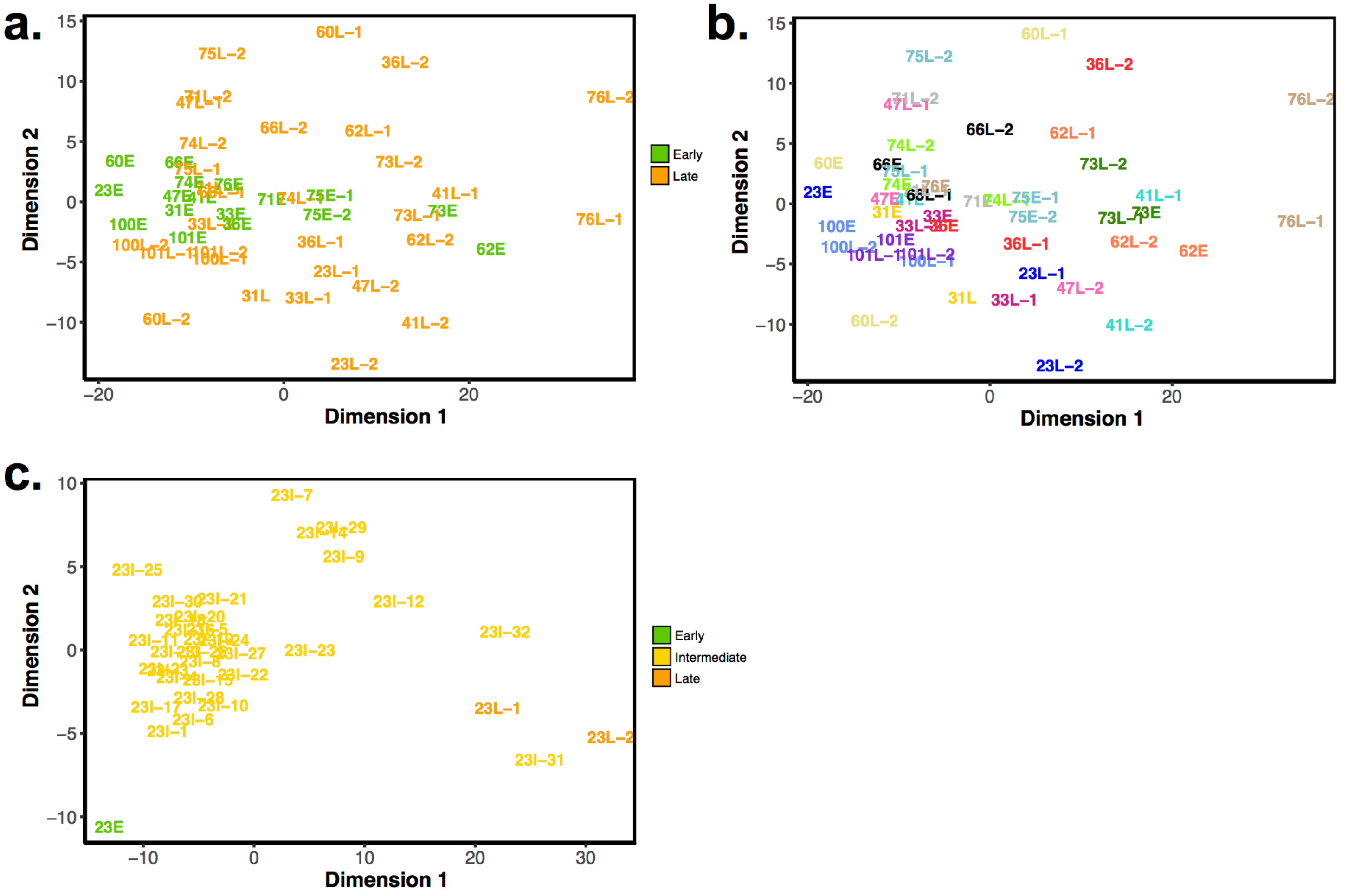

**Figure S2**. Non-metric multidimensional scaling (NMDS) plots of the truncated set of 48 *P. aeruginosa* clinical isolates depicting volatilome dissimilarity defined by Euclidean distance **(a)** colored by collection time and **(b)** colored by patient; **(c)** NMDS plot of the 35 *P. aeruginosa* clinical isolate volatilomes from Patient 23 depicting volatilome dissimilarity defined by Euclidean distance.

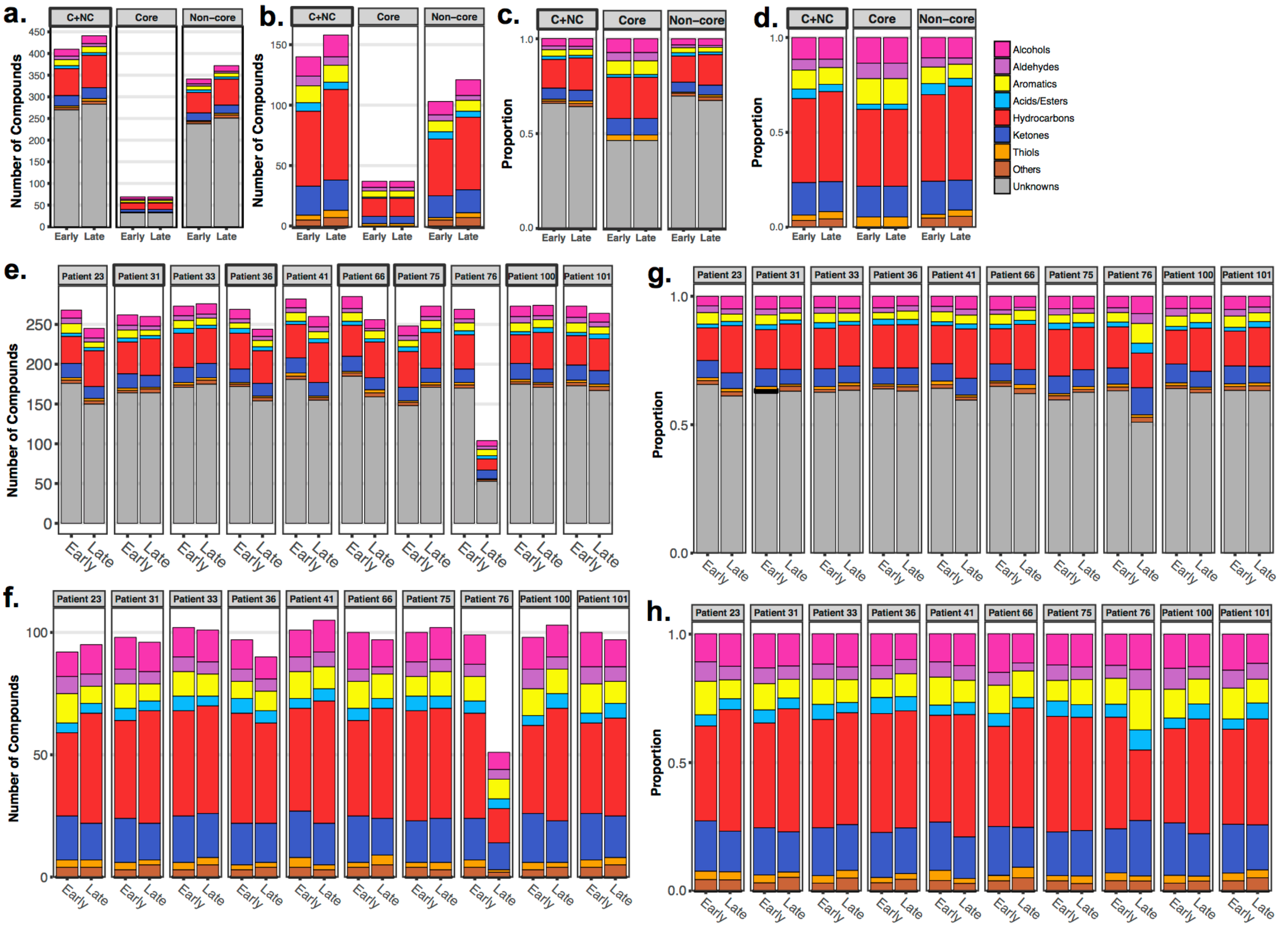

**Figure S3**. Size and chemical composition of volatile compounds metabolized by the selected ten Early-Late pairs of *P. aeruginosa* chronic infection CF isolates. **(a)** Identified compounds; **(b)** Identified and unknown compounds; **(c)** Identified compounds, scaled to 100%; **(d)** Identified and unknown compounds, scaled to 100%. Size and chemical composition of volatile compounds metabolized by each of the ten Early-Late pairs of *P. aeruginosa* chronic infection CF isolates. **(e)** Identified compounds; **(f)** Identified and unknown compounds; **(g)** Identified compounds, scaled to 100%; **(h)** Identified and unknown compounds, scaled to 100% C = Core compounds, defined as detected in 95% or more of samples; NC = Non-core compounds; C+NC = sum of Core and Non-core compounds.

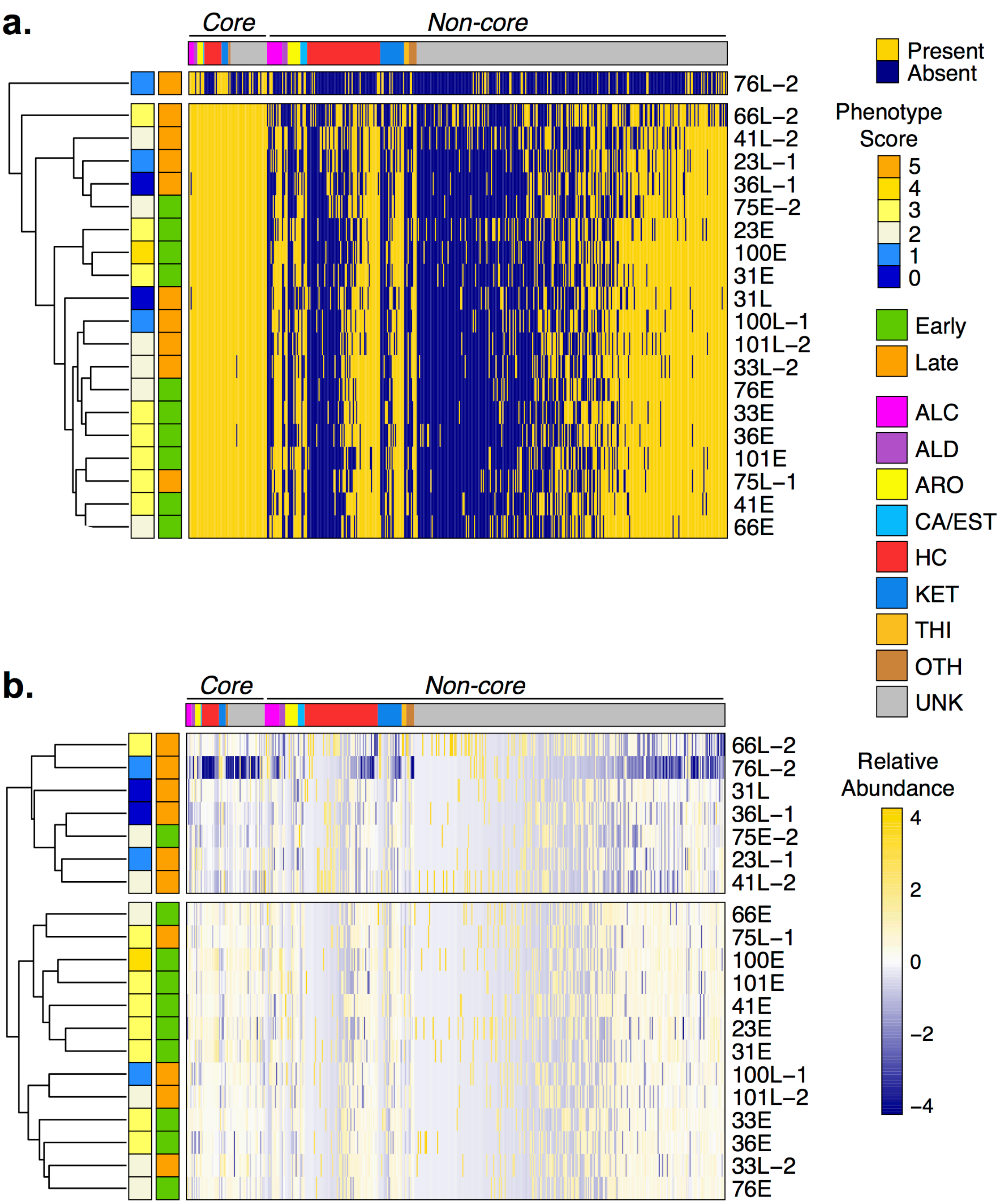

**Figure S4**. Hierarchical clustering analysis (HCA) of the ten Early-Late pairs of *P. aeruginosa* clinical CF isolates, based on **(a)** presence and absence and **(b)** relative abundance of 472 volatile compounds. Core compounds are present in at least 95% of isolates; Non-core compounds are present in less than 95%. Volatiles are in columns. Clustering is based on rows (isolates), which are color-coded by their phenotype score (left color block) and relative time of collection (right color block).

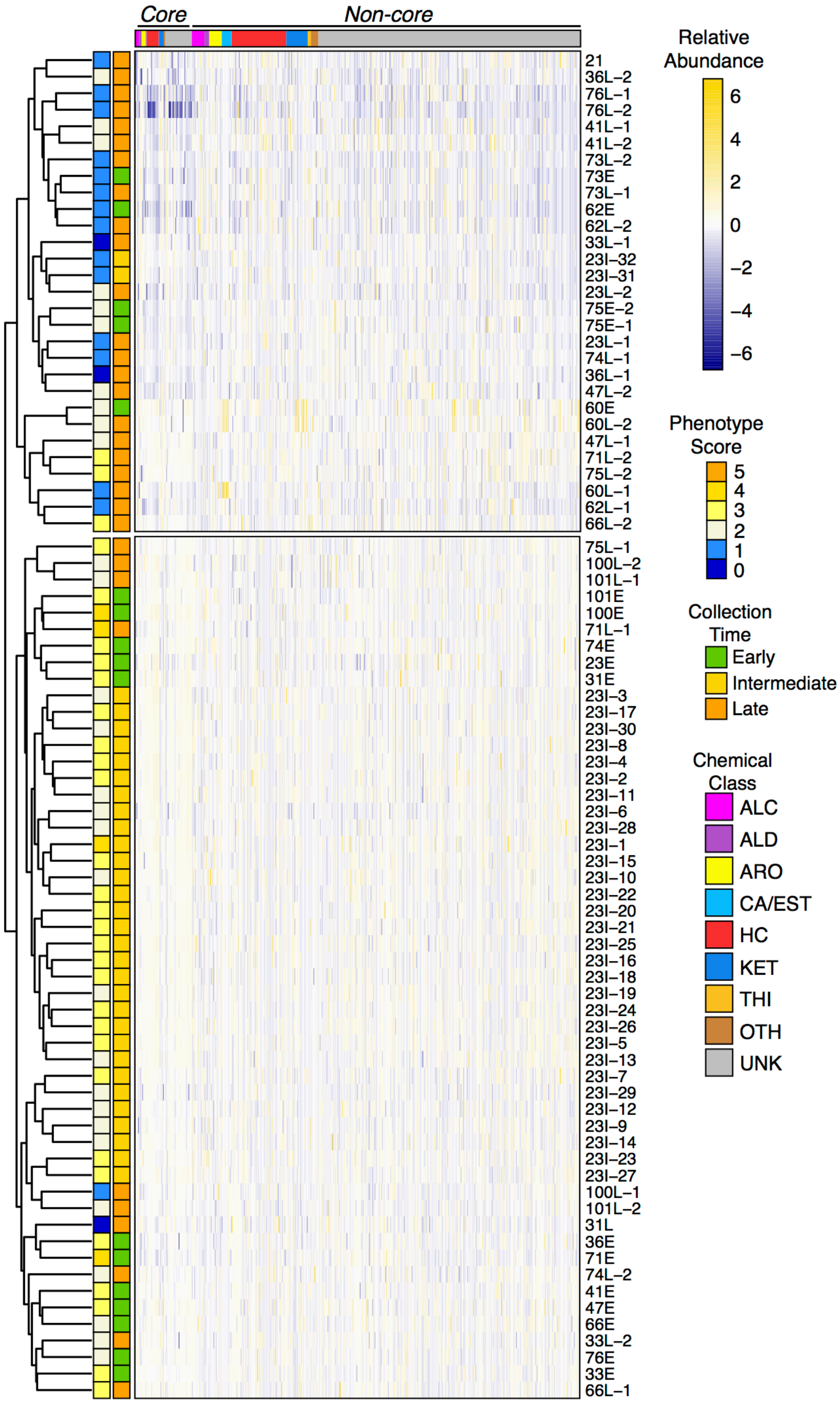

**Figure S5.** Hierarchical clustering analysis (HCA) of the 81 *P. aeruginosa* clinical CF isolates, based on the relative abundance of 539 volatile compounds. Core compounds are present in at least 95% of isolates; Non-core compounds are present in less than 95%. Volatiles are in columns (standardized relative abundance). Clustering is based on rows (isolates), which are color-coded by their phenotype score (left color block) and relative time of collection (right color block).

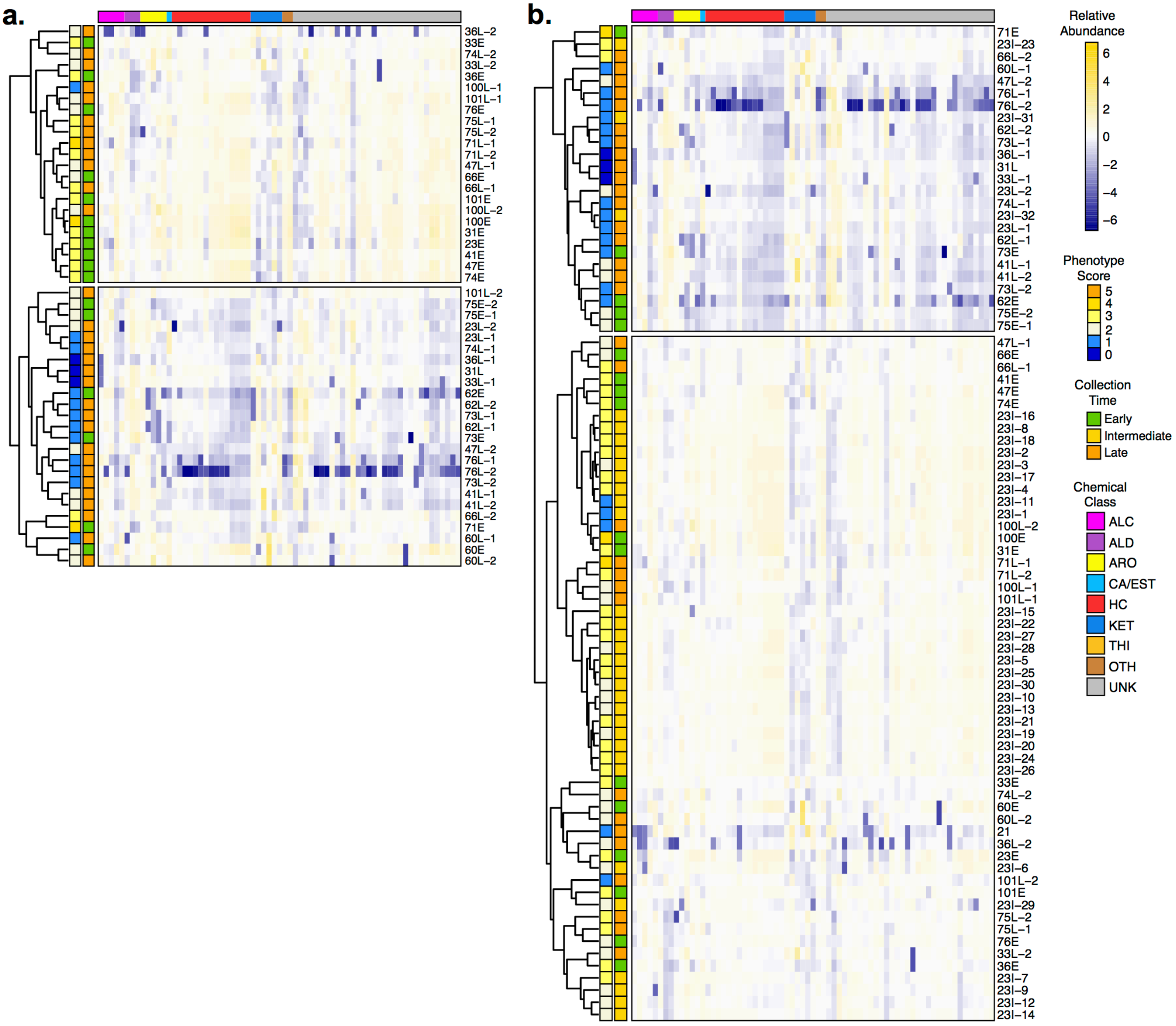

**Figure S6**. Hierarchical clustering analysis (HCA) of **(a)** the truncated set of 48 and **(b)** full set of 81 *P. aeruginosa* clinical CF isolates, based on the relative abundance of the 69 core volatile compounds. Volatiles are in columns (standardized relative abundance). Clustering is based on rows (isolates), which are color-coded by their phenotype score (left color block) and relative time of collection (right color block).

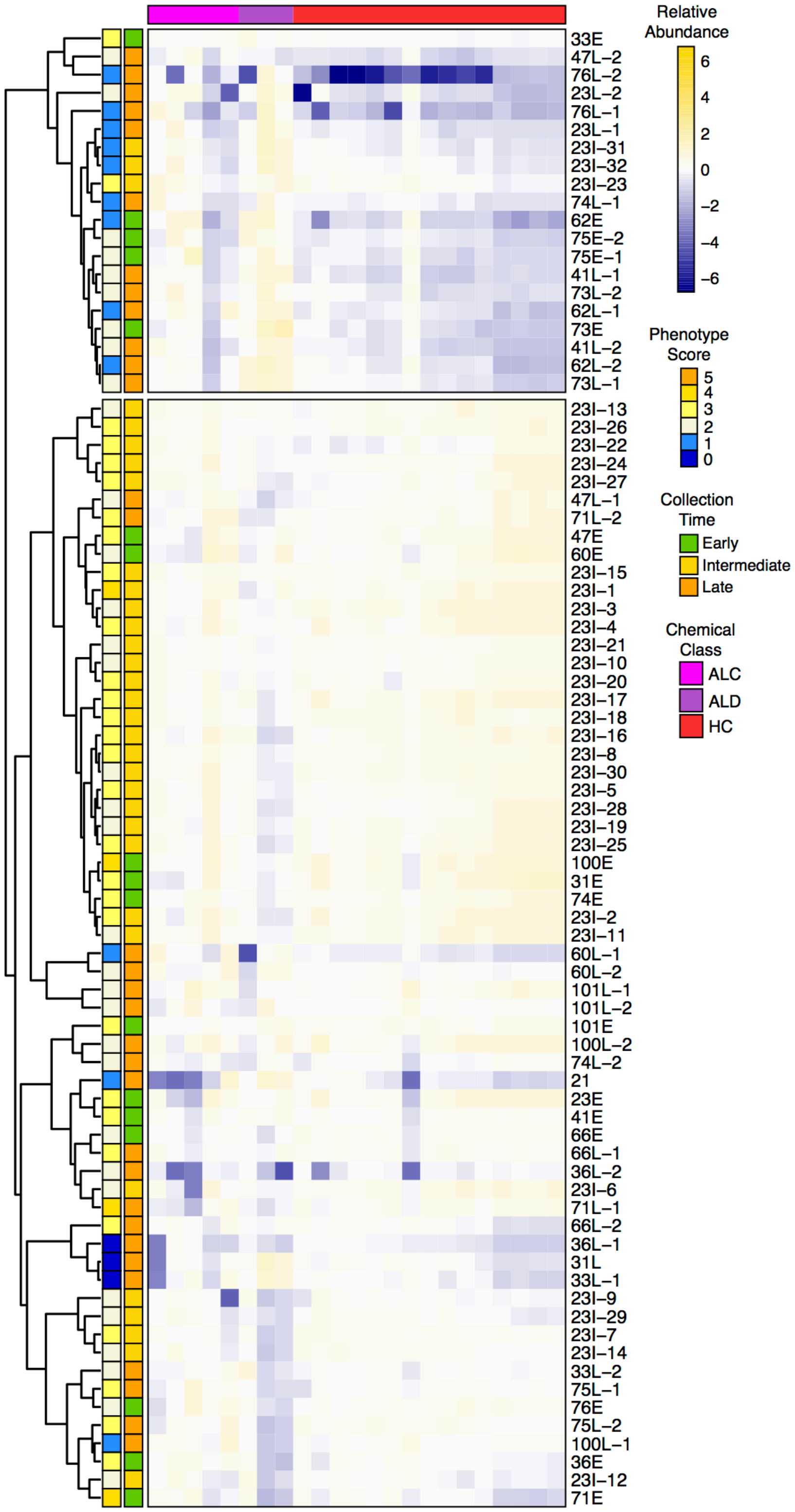

**Figure S7**. Hierarchical clustering analysis (HCA) of the 81 *P. aeruginosa* clinical CF isolates, based on the relative abundance of 23 core alcohols, aldehydes, and hydrocarbons. Volatiles are in columns (standardized relative abundance). Clustering is based on rows (isolates), which are color-coded by their phenotype score (left color block) and relative time of collection (right color block).
