## Supplementary material for "*PSEUDOMONAS AERUGINOSA* VOLATILOME CHARACTERISTICS AND ADAPTATIONS IN CHRONIC CYSTIC FIBROSIS LUNG INFECTIONS": Miscellaneous Information

**Miscellaneous Table 1.** (a) GC×GC method; (b) mass spectrometry method; (c) data processing method; (d) autosampler method.

| Supplementary Table 1a: GC×GC Method | |
|---|---|
| Instrument description | Agilent® 7890B |
| Column configuration | Column 1: Rxi®-624Sil MS, 60 m × 0.25 mm × 1.4 $\mu$m<br>Column 2: Stabilwax®, 1 m × 0.25 mm × 0.5 $\mu$m |
| Carrier gas | Helium, 2 mL·min$^{-1}$ (constant) |
| Front inlet type | Gerstel® |
| Front inlet mode | Splitless |
| Front inlet septum purge flow | 1 mL·min$^{-1}$ |
| Front inlet septum purge time | 180 s |
| Front inlet purge flow | 40 mL·min$^{-1}$ |
| Front inlet total purge flow | 42 mL·min$^{-1}$ |
| Front inlet temperature | 250 °C |
| Oven equilibration time | 5 s |
| Primary oven temperature ramp | Initial temperature: 35 °C<br>Initial time: 30 s<br>Ramp rate: 5 °C·min$^{-1}$<br>Final temperature: 230 °C<br>Hold time: 5 min |
| Secondary oven temperature offset | 5 °C (relative to primary oven) |
| Modulator temperature offset | 20 °C (relative to secondary oven) |
| Modulation timing | Modulation period: 2.00 s<br>Hot pulse time: 0.50 s<br>Cold pulse time: 0.50 s |
| Transfer line temperature | 250 °C |

| Supplementary Table 1b: Mass Spectrometry Method | |
|---|---|
| Instrument description | LECO® Pegasus® 4D |
| Use GC method total time for MS method total time | Yes |
| Acquisition delay | 180 s |
| Filament active time | 180 s to end of run |
| Start mass/End mass | 35/400 |
| Acquisition rate | 100 spectra·s$^{-1}$ |
| Optimized voltage offset | +50 V |
| Electron energy | -70 eV |
| Ion source temperature | 200 °C |
| Acquisition masses | TIC |

| Supplementary Table 1c: Data Processing Method | |
|---|---|
| Software description | LECO® ChromaTOF® (version 4.51) |
| Baseline tracking/Offset | Entire run/0.5 (through middle of noise) |

| | |
|---|---|
| Data points for smoothing | Auto |
| First dimension peak width | 12 |
| Mass spectral match required to combine peaks | 600 |
| Second dimension peak width | 0.15 |
| Minimum subpeak signal-to-noise (S/N) | 6 |
| Integration approach | Traditional |
| Processing | S/N: 50<br>Number of apexing masses: 2 |
| Library identity search mode/Library search mode | Normal/Forward |
| Masses to library search | All |
| Minimum/Maximum molecular weight | 35/550 |
| Mass threshold | 10 |
| Libraries for searching | NIST® 2011 |
| Mass to use for area/height calculation | Unique mass |
| Analyte match criteria | Spectral match mass threshold: 10<br>Minimum spectral similarity match: 600<br>Max. number of modulation periods apart: 3<br>Max. retention time difference: ±0.2<br>S/N for second peak find: 5 |
| Criteria for keeping analytes | Min. number of samples that contain analyte: 1 |

| **Supplementary Table 1d: Autosampler Method** | |
|---|---|
| Instrument description | Gerstel® MPS Pro® |
| Software description | Gerstel® Maestro® (version 1.5.3.2) |
| *Sampling Parameters* | |
| Cooled tray temperature | 4 °C |
| Solid-phase microextraction (SPME) | Manufacturer: Supelco®<br>Fiber type: PDMS/CAR/DVB (2 cm) |
| Incubation time | 5 min |
| Agitator parameters | Temperature: 50 °C<br>On time: 10 s<br>Off time: 1 s<br>Speed: 600 RPM |
| Vial penetration | 21 mm |
| Extraction time | 10 min |
| Injection penetration | 67 mm |
| Desorption time | 180 s |

| *Inlet (CIS) Parameters* | |
|---|---|
| Initial temperature | 250 °C |
| Equilibrium time | 0.05 min |

| | |
|---|---|
| Initial time | 0.10 min |
| Ramp rate | 12 °C·$s^{-1}$ |
| End temperature | 250 °C |
| Hold time | 10.5 min |

**Miscellaneous Table 2.** List of removed peaks, which include poorly modulated chromatographic features, such as atmospheric gasses, and common contaminants and artifacts that are associated with the SPME or GC stationary phase polymer decomposition (e.g., siloxanes and oximes), identified based on comparisons to system blanks, with removed peak names assigned using a NIST MS match score >600.

| Identification (based on mass spectral match of ≥ 600/999) |
|---|
| 1-(1-bromoethyl)-1,3-dioxolane |
| 1-[2,4-Bis(trimethylsiloxy)phenyl]-2-[(4-trimethylsiloxy)phenyl]propan-1-one |
| 1,1,1,3,5,5,5-heptamethyltrisiloxane |
| 1,1,1,3,5,7,7,7-octamethyl-3,5-bis(trimethylsiloxy)tetrasiloxane |
| 1,1,1,5,5,5-hexamethyl-3,3-bis[(trimethylsilyl)oxy]-trisiloxane |
| 1,3-diisopropoxy-1,3-dimethyl-1,3-disilacyclobutane |
| 1,3-dioxolane |
| 1,3-dioxolane-2-methanol |
| 16-methyl-heptadecane-1,2-diol, trimethylsilyl ether |
| 2-(1-bromoethyl)-1,3-dioxolane |
| 2-(3-bromo-5,5,5-trichloro-2,2-dimethylpentyl)-1,3-dioxolane |
| 2-(6-heptynyl)-1,3-dioxolane |
| 2-[(tert-butyldimethylsilyl)oxy]-butanedioic acid, bis(tert-butyldimethylsilyl) ester |
| 2-[(trimethylsilyl)oxy] benzoic acid, trimethylsilyl ester |
| 2-[(trimethylsilyl)oxy]-ethanol |
| 2-butyl-1,3-dioxolane |
| 2-heptyl-1,3-dioxolane |
| 2-hydroxy-5-methoxybenzoic acid, tert-butyldimethylsilyl ether, tert-butyldimethylsilyl ester |
| 2-pentadecyl-1,3-dioxolane |
| 2,5-bis[(trimethylsilyl)oxy]-benzaldehyde |
| 2',6'-Dihydroxyacetophenone, bis(trimethylsilyl) ether |
| 3-hydroxy-4-methoxybenzaldehyde, tert-butyldimethylsilyl ether |
| 3,3,6,6-tetramethyl-1,2,4,5-tetroxane |
| 3,5-diethoxy-1,1,1,7,7,7-hexamethyl-3,5-bis(trimethylsiloxy)-tetrasiloxane |
| 5-trimethylsilylmethyl-1-trimethylsilyl-spiro[2.4]hept-5-ene |
| 7,7'-dihydro-6,6'-bis(trimethylsilyl)methylheptalene |
| N,O-bis(tert-butyldimethylsilyl)-2-amino-4-nitrophenol |
| Argon |
| Arsenous acid, tris(trimethylsilyl) ester |
| bis(trimethylsilyl)-2-methyl-2-(p-methoxy)-mandalate |
| bis[(trimethylsilyl)oxy]phosphinyl] acetic acid, trimethylsilyl ester |
| Boron trifluoride |
| Butyltrimethylsilane |
| Carbon dioxide |
| Difluorodimethylsilane |
| Dimethylsilanediol |
| Dodecamethylcyclohexasiloxane |
| Hexamethylcyclotrisiloxane |
| Hexamethyldisiloxane |
| Hydrogen azide |
| Methanephosphonofluoridic acid trimethylsilyl ester |
| Methanephosphonofluoridic acid, fluoroanhydride, tert-butyldimethylsilyl ester |

| |
|---|
| Methoxyphenyloxime |
| Nickel tetracarbonyl |
| Nitrous oxide |
| Octamethylcyclotetrasiloxane |
| Oxalic acid, 2-ethylhexyl hexyl ester |
| Oxalic acid, 6-ethyloct-3-yl isohexyl ester |
| Oxalic acid, allyl decyl ester |
| Oxalic acid, allyl hexadecyl ester |
| Oxalic acid, allyl isohexyl ester |
| Oxalic acid, allyl nonyl ester |
| Oxalic acid, allyl octadecyl ester |
| Oxalic acid, allyl octyl ester |
| Oxalic acid, allyl tridecyl ester |
| Oxalic acid, butyl isobutyl ester |
| Oxalic acid, cyclobutyl decyl ester |
| Oxalic acid, cyclobutyl octadecyl ester |
| Oxalic acid, cyclobutyl tetradecyl ester |
| Oxalic acid, cyclohexyl tetradecyl ester |
| Oxalic acid, isobutyl nonyl ester |
| Oxalic acid, isobutyl pentyl ester |
| Oxalic acid, isohexyl pentyl ester |
| Oxalic acid, tetradecyl ester |
| Propanoic acid, dimethyl(isopropyl)silyl ester |
| Pyrocatechol, bis(tert-butyldimethylsilyl) ether |
| Tetramethylsilane |
| trans-Trimethyl-3-penten-2-ylsilane |
| Trichlorodocosylsilane |
| Trimethyl[2-methylene-1-(4-pentenyl)cyclopropyl]-silane |
| Trimethylsilanol |
| ***Poorly modulated peaks and/or peaks eluting prior to 358 s*** |
| 1-tetrazol-2-ylethanone |
| 1,2-propanediamine |
| 11-amino-dodecanoic acid, methyl ester |
| 2,6-difluoro-3-methylbenzoic acid, 2,3-dichlorophenyl ester |
| 3,3,4,4-tetrafluorohexane |
| N,N-dimethylmethylamine |
| Chloromethane |
| Chloromethanesulfonyl chloride |
| Fluoroethyne |
| Phenylephrine |
| Tuaminoheptane |

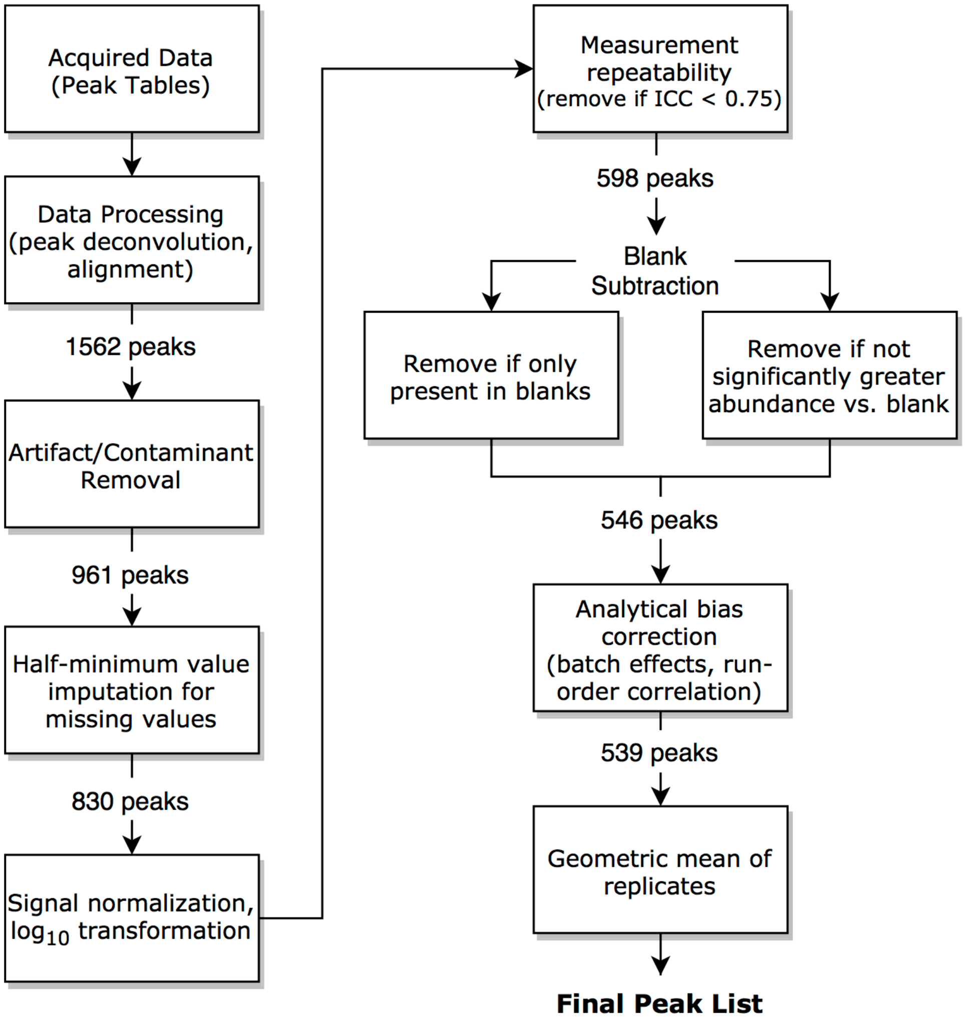

**Miscellaneous Figure 1**. Flow chart of data processing.

**Miscellaneous Figure 7**. Representative GC×GC chromatograms of the selected ten Early-Late pairs of *P. aeruginosa* clinical CF isolates (parts **(a)** – **(j)**). Dark blue represents the baseline, and peak intensity is depicted using a color gradient from light blue to dark red. Chromatographic regions of $^{1}t_{R}$ < 200 s and $^{2}t_{R}$ < 0.5 s were excluded for visual clarity.

#### (a) Isolates 23E (top) and 23L-1 (bottom)

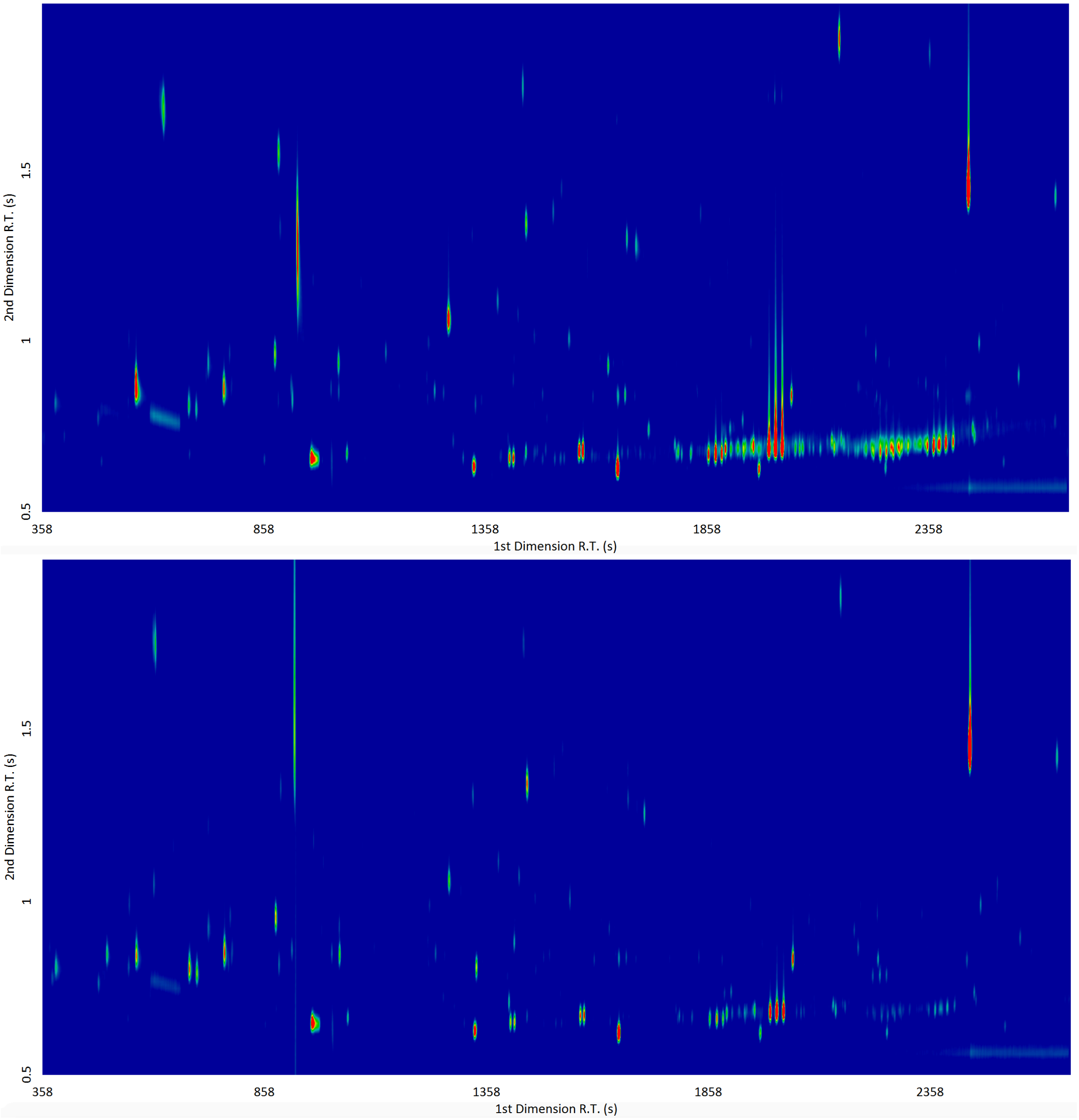

### (b) Isolates 31E (top) and 31L (bottom)

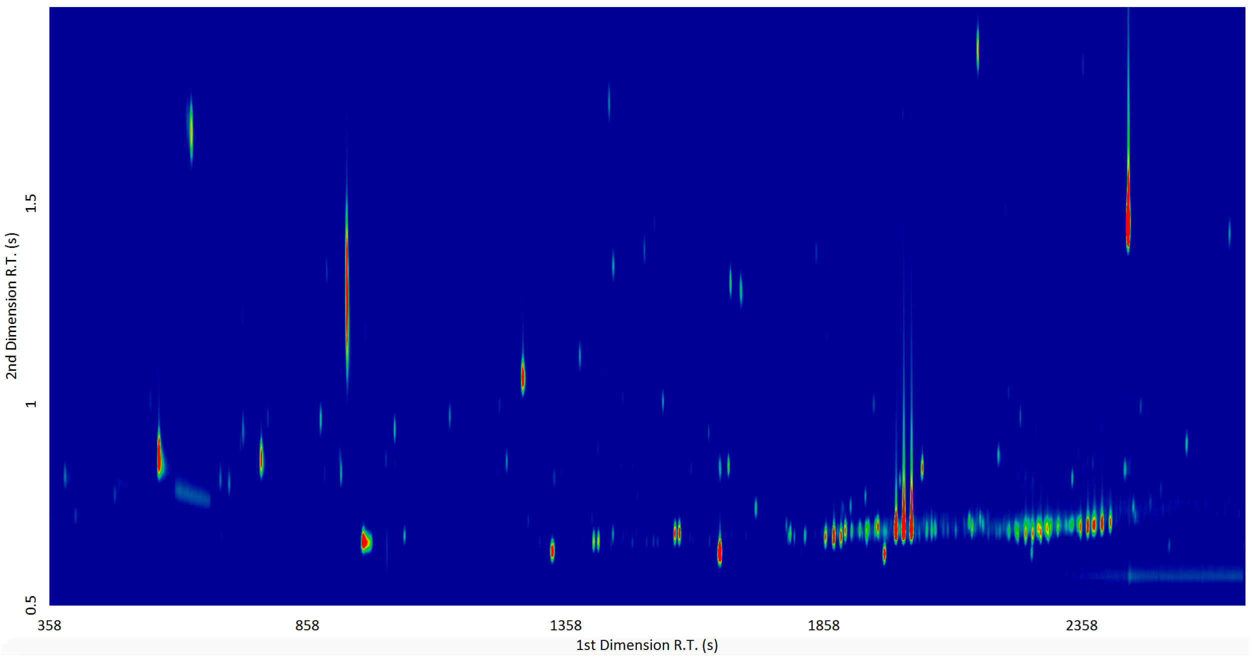

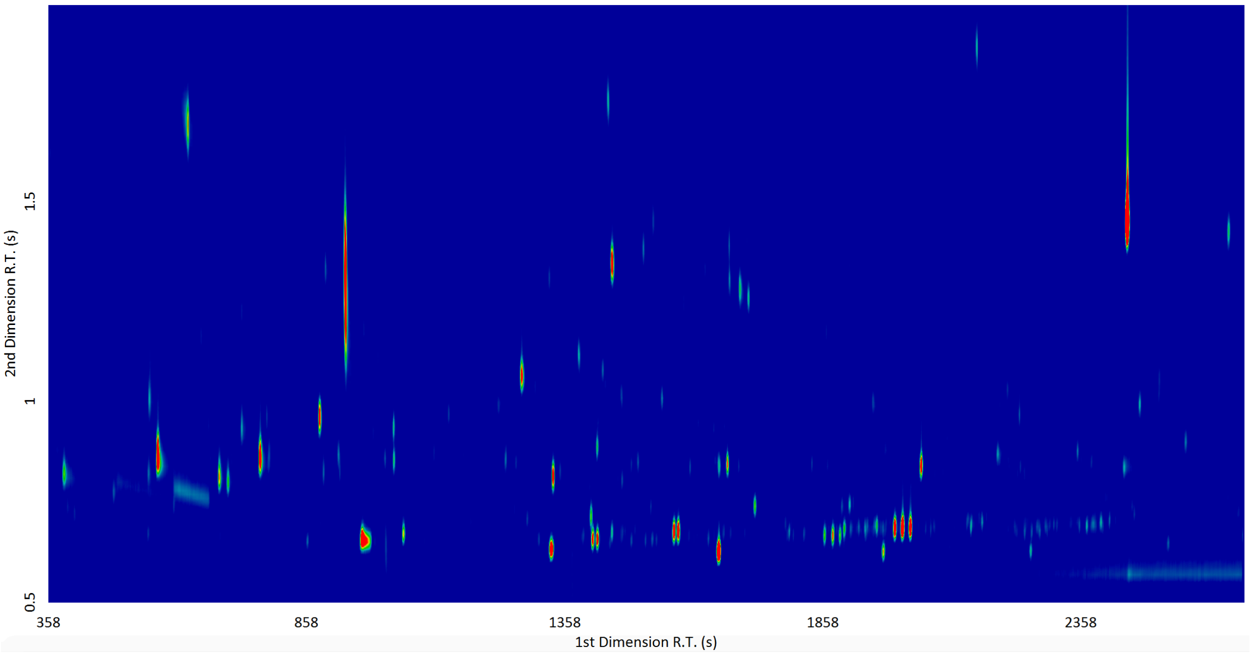

### (c) Isolates 33E (top) and 33L-2 (bottom)

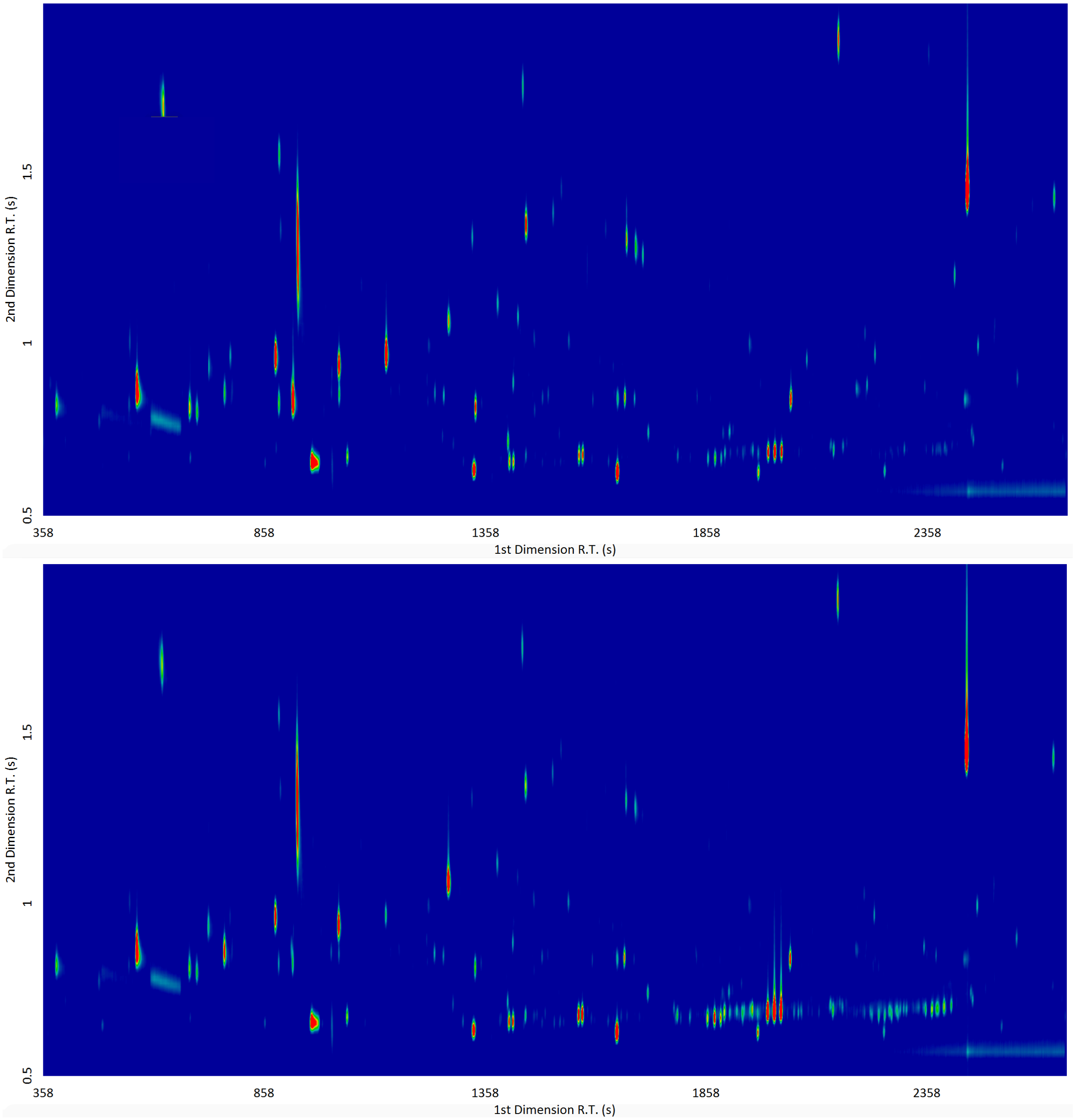

**(d) Isolates 36E (top) and 36L-1 (bottom)**

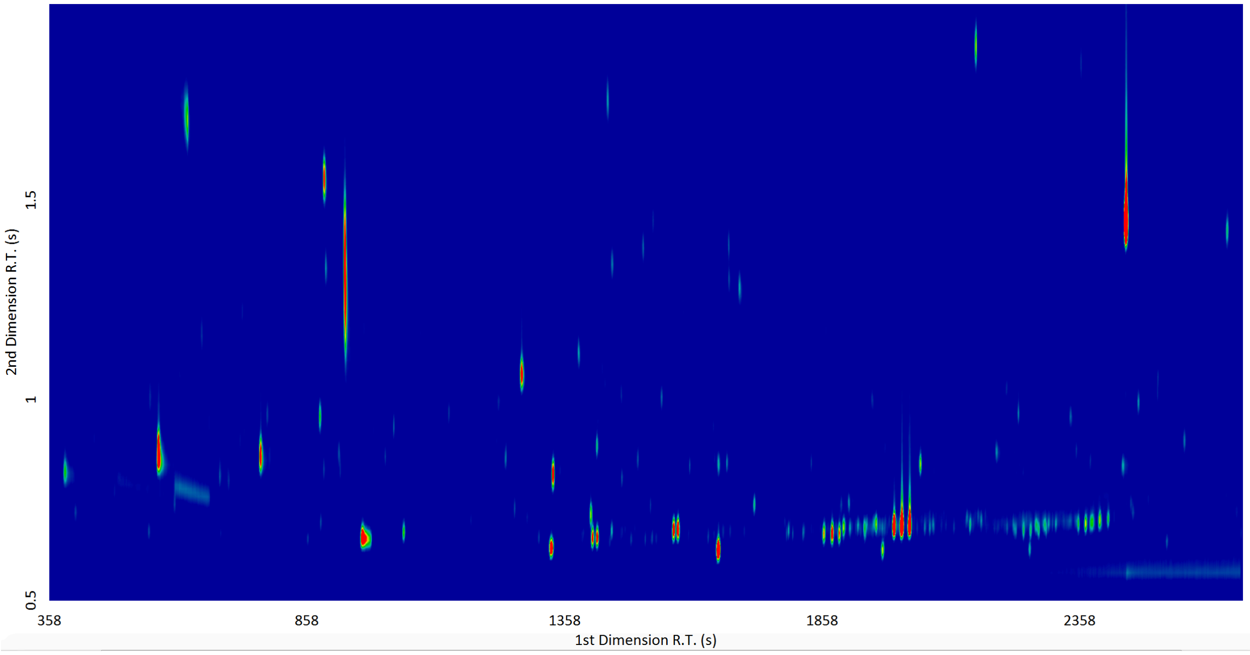

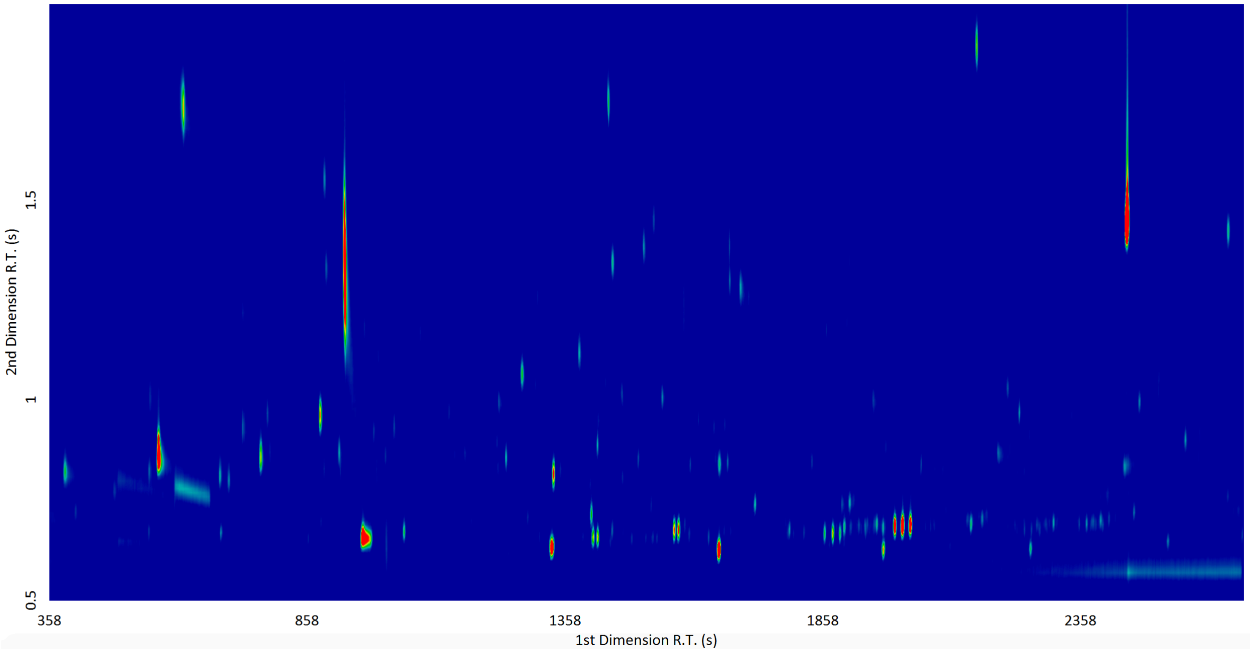

### (e) Isolates 41E (top) and 41L-2 (bottom)

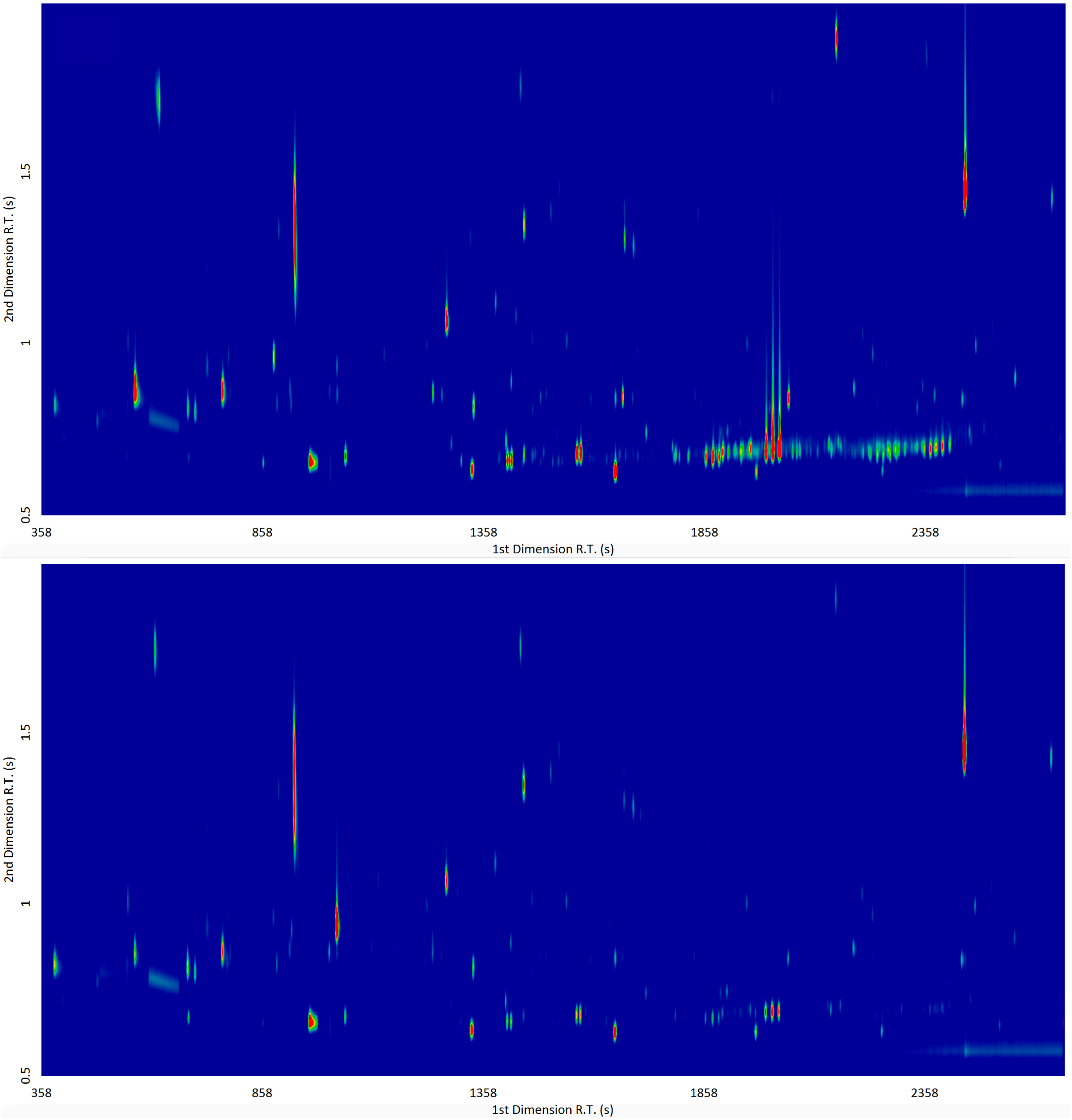

### (f) Isolates 66E (top) and 66L-1 (bottom)

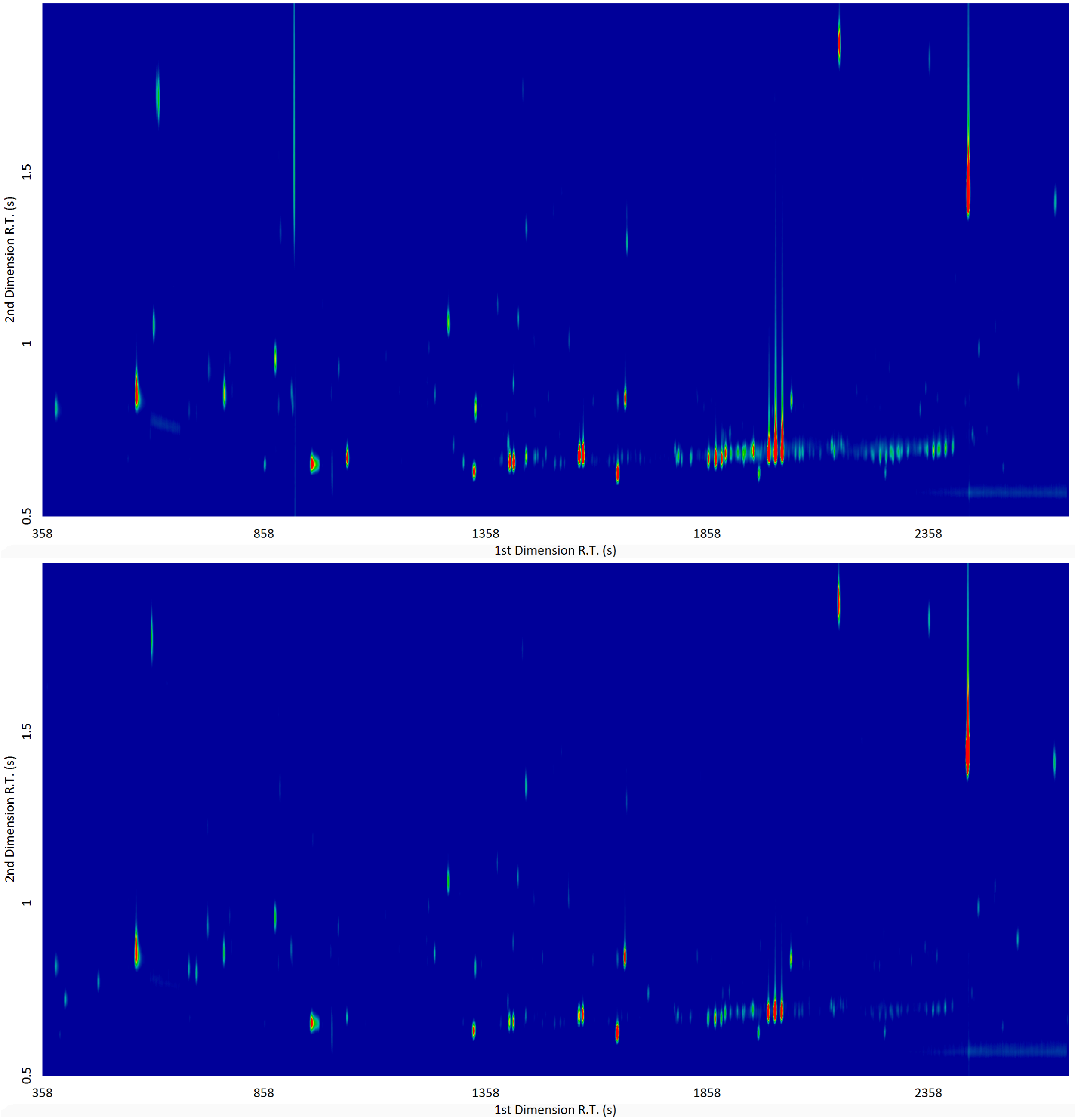

### (g) Isolates 75E-1 (top) and 75L-1 (bottom)

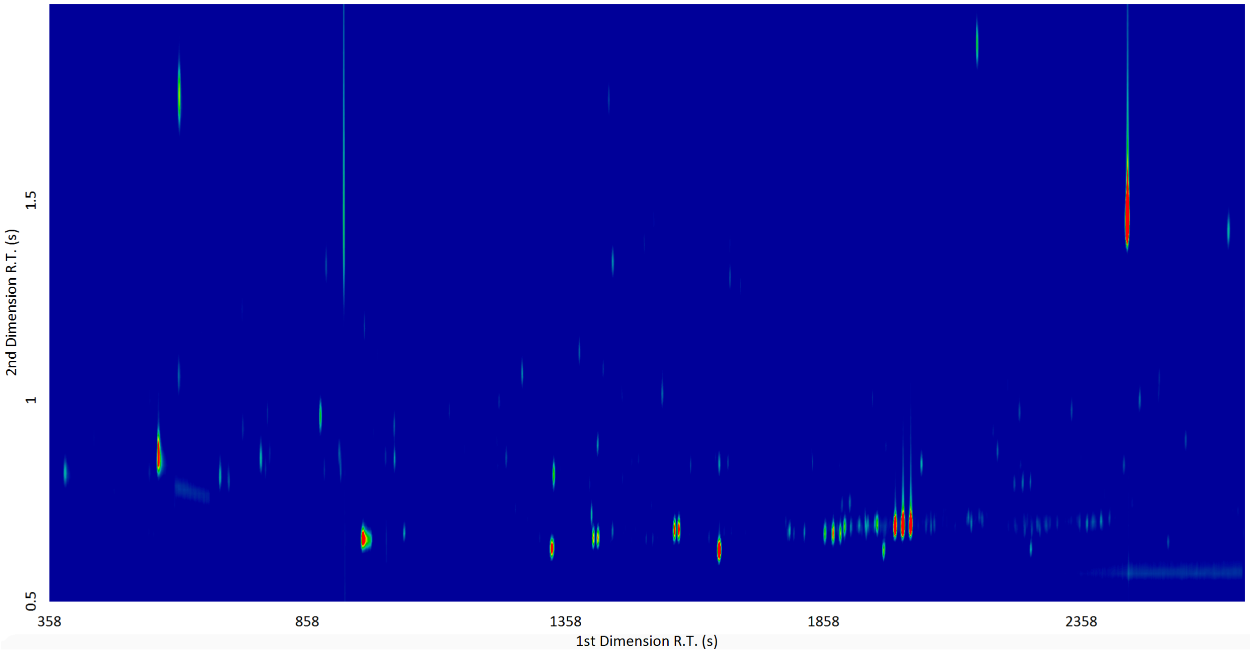

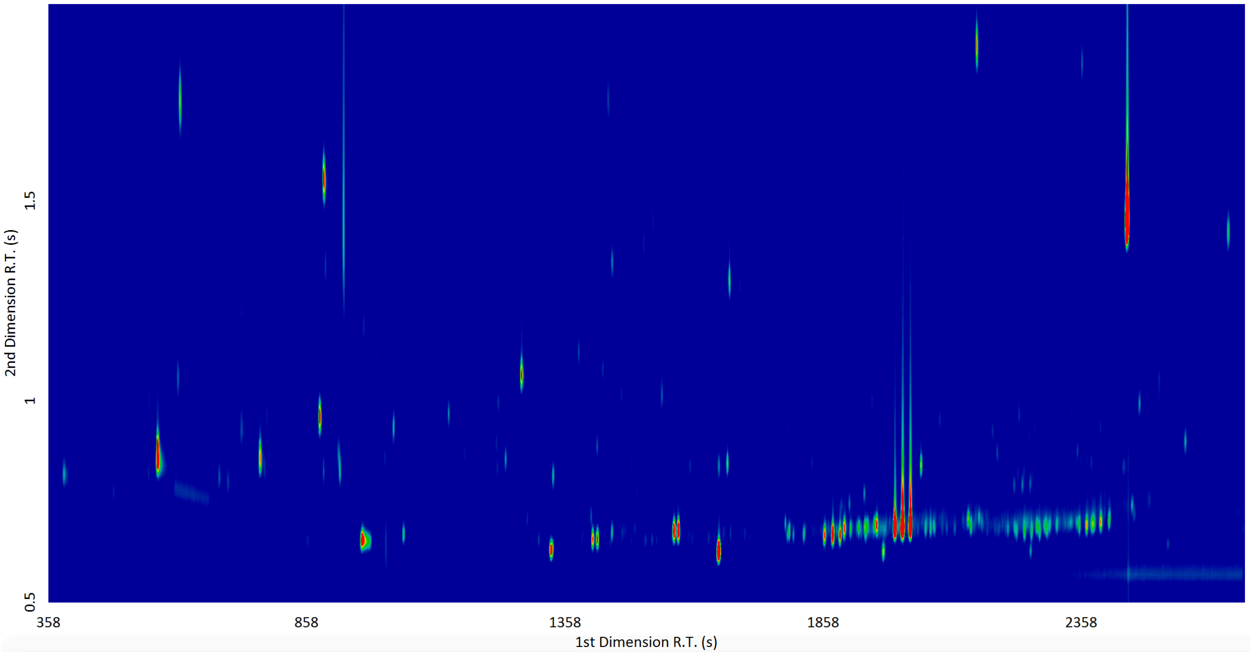

**(h) Isolates 76E (top) and 76L-2 (bottom)**

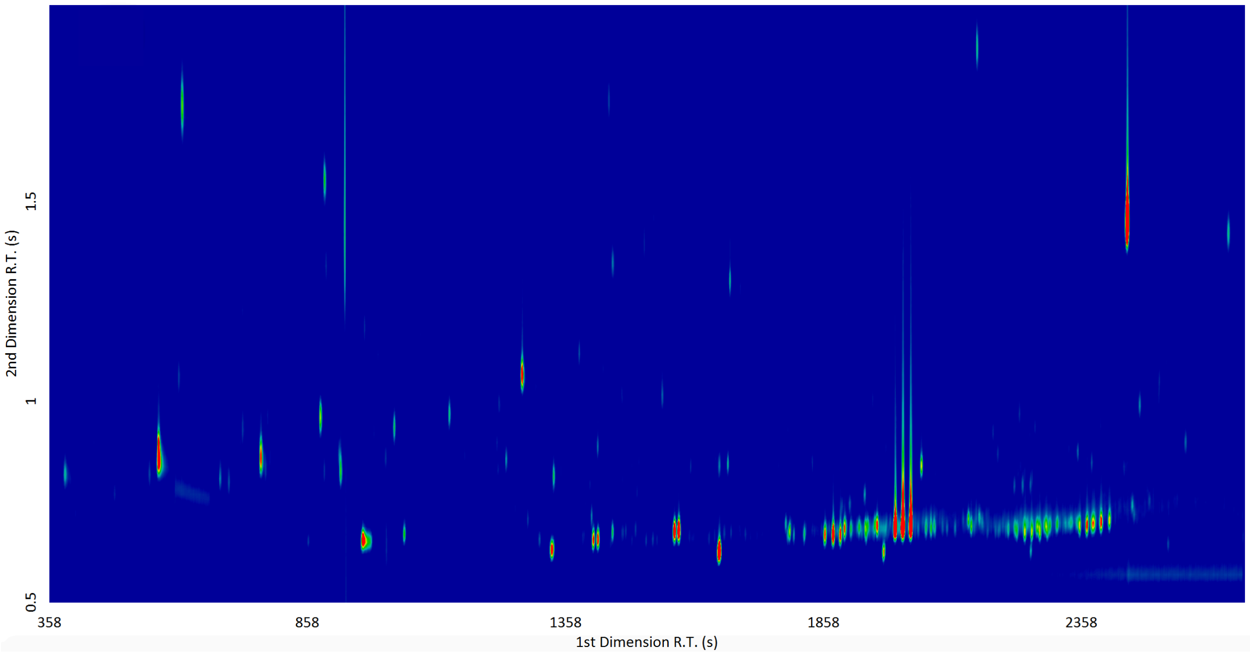

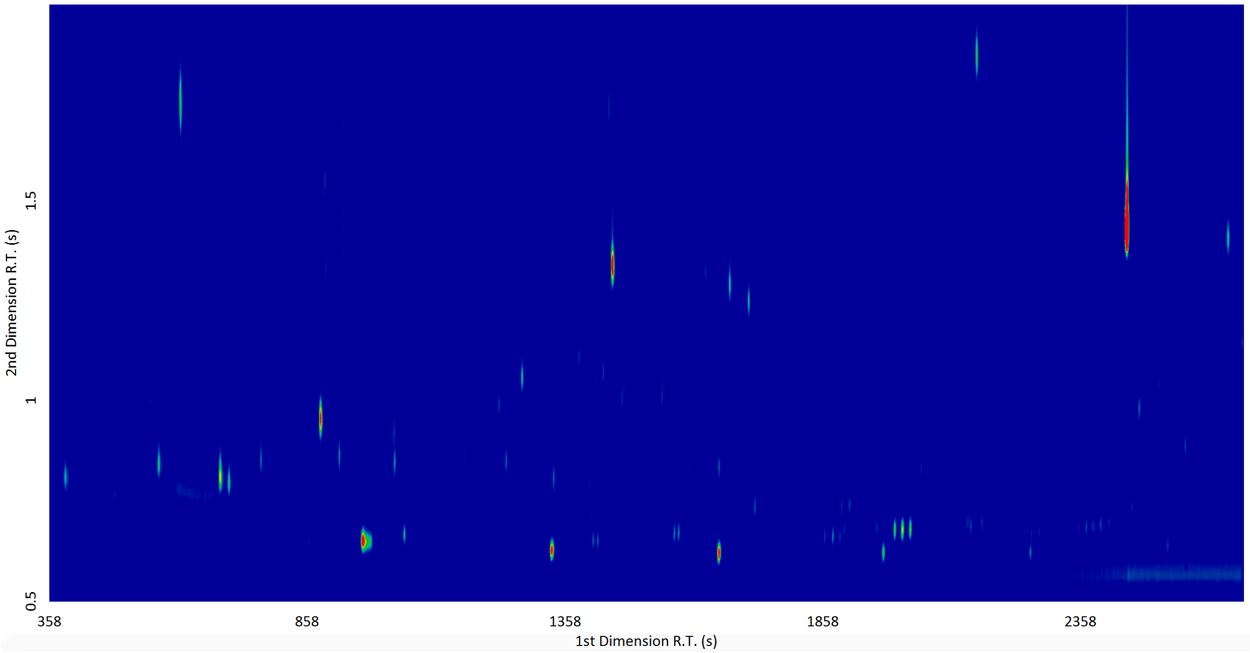

### (i) Isolates 100E (top) and 100L-1 (bottom)

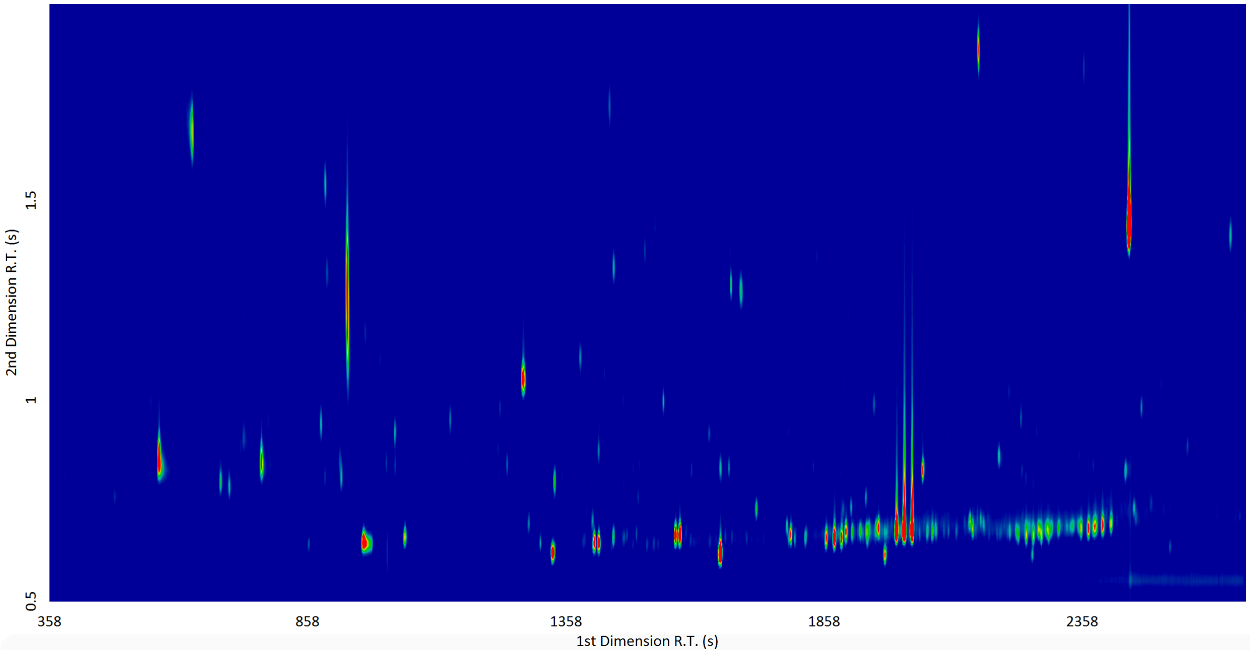

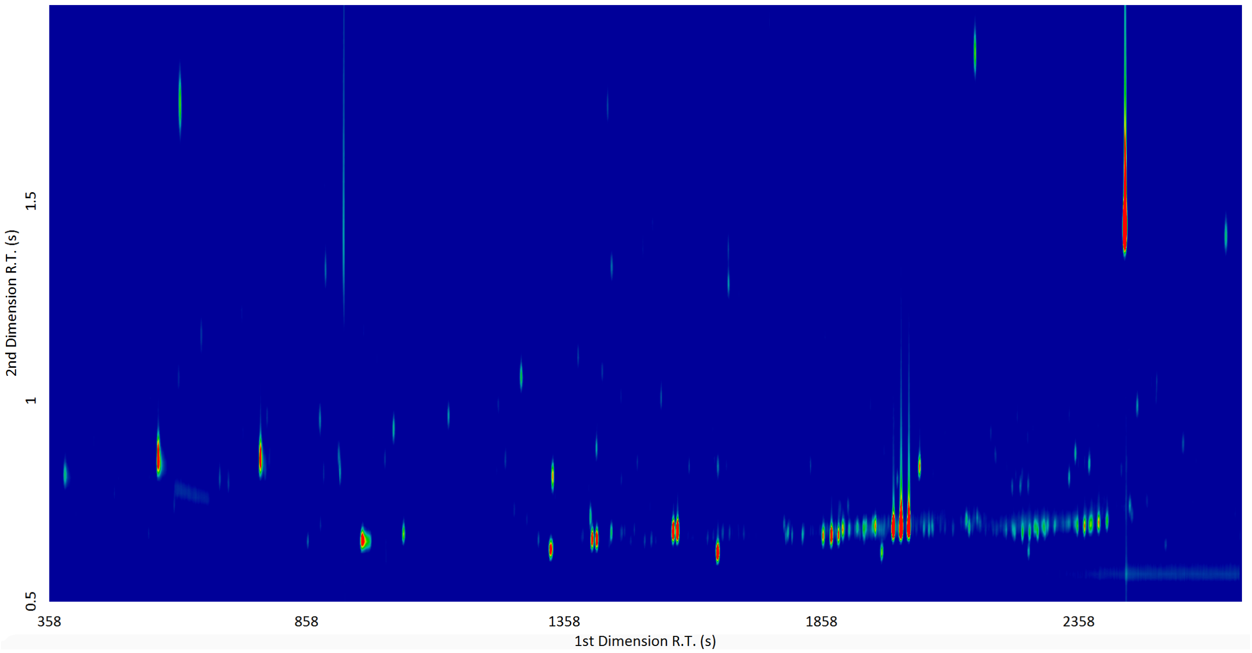

### (j) Isolates 101E (top) and 101L-2 (bottom)

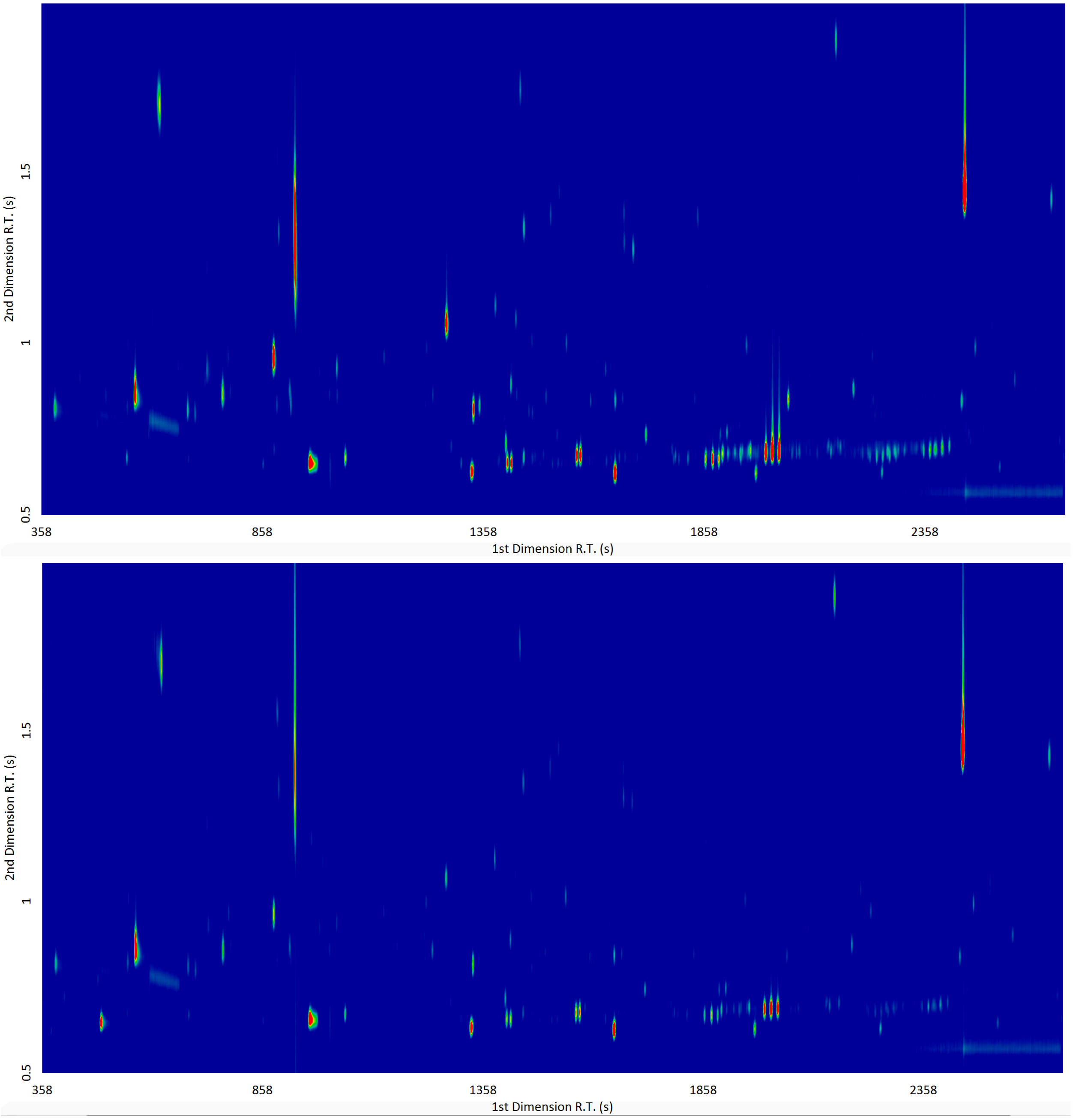
